# G9a methyltransferase governs cell identity in the lung and is required for KRAS G12D tumor development and propagation

**DOI:** 10.1101/2020.04.20.050328

**Authors:** Ariel Pribluda, Anneleen Daemen, Anthony Lima, Xi Wang, Marc Hafner, Chungkee Poon, Zora Modrusan, Anand Kumar Katakam, Oded Foreman, Jeffrey Eastham, Jefferey Hung, Benjamin Haley, Julia T Garcia, Erica L. Jackson, Melissa R. Junttila

## Abstract

Lung development, integrity and repair rely on precise Wnt signaling, which is corrupted in diverse diseases, including cancer. Here, we discover that G9a methyltransferase regulates Wnt signaling in the lung by controlling the transcriptional activity of chromatin-bound β-catenin, through a non-histone substrate. Inhibition of G9a induces transcriptional, morphologic, and molecular changes consistent with alveolar type 2 (AT2) lineage commitment. Mechanistically, G9a activity functions to support regenerative properties of KrasG12D tumors and normal AT2 cells – the predominant cell of origin of this cancer. Consequently, G9a inhibition prevents *KrasG12D* lung adenocarcinoma tumor formation and propagation,and disrupts normal AT2 cell trans-differentiation. Consistent with these findings, low G9a expression in human lung adenocarcinoma correlates with enhanced AT2 gene expression and improved prognosis. These data reveal G9a as a critical regulator of Wnt signaling, implicating G9a as a potential target in lung cancer and other AT2-mediated lung pathologies.

## Introduction

Lung cancer is the leading cause of cancer mortality in men and women, surpassing combined deaths from colon, prostate and breast cancer (www.cancer.org, 2019). Approximately 40% of non-small cell carcinoma (NSCLC) is of the adenocarcinoma subtype. Genomic alterations of KRAS and TP53 mutations are the most frequent events within this tumor subtype (Cancer Genome Atlas Research, 2014). Considerable evidence indicates that alveolar type 2 (AT2) cells are the predominant cell of origin of lung adenocarcinoma (LUAD) (Desai et al., 2014; Mainardi et al., 2014; Sutherland et al., 2014; Xu et al., 2012).

Sustained tumor growth is maintained, in some instances, by a subset of plastic, stem-like cells, referred to as tumor-propagating cells (TPCs) (Batlle and Clevers, 2017; Beck and Blanpain, 2013). This rare cell subset is less sensitive to therapeutic intervention, and functionally contributes to tumor re-growth following treatment responses (Beck and Blanpain, 2013). TPCs are hypothesized to evolve extensively from the tumor-initiating event (Reya et al., 2001; Shackleton et al., 2009), yet the fidelity to its cell of origin remain undefined. TPCs in LUAD have been previously characterized and were shown to be required for KrasG12D tumor self-renewal (Zheng et al., 2013). Moreover, a TPC gene signature correlates with poor prognosis in non-small cell lung adenocarcinoma patients (Zheng et al., 2013). While these data implicate a critical function for TPCs in driving tumor progression, it is unknown how these rare cells maintain their stem-like properties.

Wnt signaling is of critical importance in the lung, controlling tissue development, homeostasis and repair processes following lung damage. Wnt signaling is critical in the distal airway to maintain AT2 cell fate and function in the alveolar compartment. A subset of AT2 cells serve as adult tissue stem cells, replenishing themselves, as well as alveolar type 1 (AT1) cells, which are both required to maintain proper alveolar function. Deletion of β-catenin in AT2 cells leads to AT1 cell fate transdifferentiation, demonstrating that β-catenin-mediated Wnt signaling is required for AT2 cell identity (Frank et al., 2016; Nabhan et al., 2018). Along the same lines, genetic models have also shown the forced expression of a stabilized form of β-catenin engenders aberrant cell fate in mouse lung (Pacheco-Pinedo et al., 2011). Together these data, indicate that precise Wnt pathway activity is important for alveolar cell function, as well as, cell fate decisions. Furthermore, disruption of Wnt signaling equilibrium can have major consequences in some disease pathologies, including lung cancer (Nabhan et al., 2018; Tammela et al., 2017; Zacharias et al., 2018). What remains unclear is how Wnt signals are intrinsically regulated in cells to maintain or acquire facultative stem-like properties in both homeostatic and disease contexts. Understanding the mechanisms that underlie cell fate decisions may enable therapeutic approaches to enhance or prevent these processes for patient benefit.

In many biological contexts, cellular self-renewal and lineage fate commitment is controlled by epigenetic mechanisms, such as chromatin remodeling (Easwaran et al., 2014; Hemberger et al., 2009; Widschwendter et al., 2007). Recent work identified expression of the G9a lysine methyltransferase (EHMT2) as a poor prognostic factor in LUAD (Chen et al., 2010; Huang et al., 2017). Interestingly, G9a activity has been implicated in the regulation of cell identity during development primarily through its epigenetic histone methyltransferase activity (Chen et al., 2012; Epsztejn-Litman et al., 2008). While G9a methyltransferase activity and its regulation of chromatin is well-described; the non-histone targets of epigenetic regulators are emerging as critical signaling mediators. Here, we investigated how G9a methyltransferase functions to regulate cell fate in distinct cellular contexts relevant for lung tumor development, propagation and AT2 biology. We discover a mechanism of cell intrinsic Wnt signaling control that is governed by G9a activity, establishing G9a as a crucial arbitrator of cell fate gene expression in the lung. By discovering this alternative cell-intrinsic mechanism of governing Wnt signaling in cells, we are able to manipulate cell fate decisions to predictably limit the functionality of these cells to disable tumor propagation, development and AT2 trans-differentiation.

## Results

### G9a activity is required for *Kras*^*G12D*^;*Trp53* (KP) tumorsphere self-renewal

Given the association of G9a expression and poor prognosis in LUAD (Chen et al., 2010; Huang et al., 2017), we sought to examine the expression and function of G9a in primary murine *KrasG12D;p53*^−/−^ (KP) tumors. Previous work established that KP tumor self-renewal was dependent on a TPC subset. Therefore, by utilizing the previously characterized surface markers CD24, ITGB4 and NOTCH (Zheng et al., 2013), we sorted the TPC population and evaluated G9a protein expression. Using two distinct detection methods we observed a consistent and robust increase in G9a protein expression (Figure 1A-C). Next, we evaluated the requirement of G9a in self-renewal function of TPCs using *ex vivo* KP-derived organotypic cultures (ie. tumorspheres), an established surrogate for measuring the *in vivo* regenerative capability of TPCs (Zheng et al., 2013). Pharmacologic inhibition of G9a, using UNC0642 (Liu et al., 2013), resulted in a dose-dependent decrease of primary *ex vivo* KP*-*derived tumorsphere formation (Figure 1D). Pharmacologic inhibition of G9a in established tumorspheres led to a marked reduction in both histone H3 lysine 9 di- and tri-methylation marks (H3K9me2/3), consistent with potent G9a inhibition (Collins and Cheng, 2010; Epsztejn-Litman et al., 2008; Shinkai and Tachibana, 2011) (Figure 1E–figure supplement 1). Both UNC0642 treatment or short hairpin RNA (shRNA)-mediated depletion of G9a similarly impaired secondary sphere formation (Figure 1F-I), establishing a requirement for G9a activity in TPC self-renewal.

**Figure 1.**
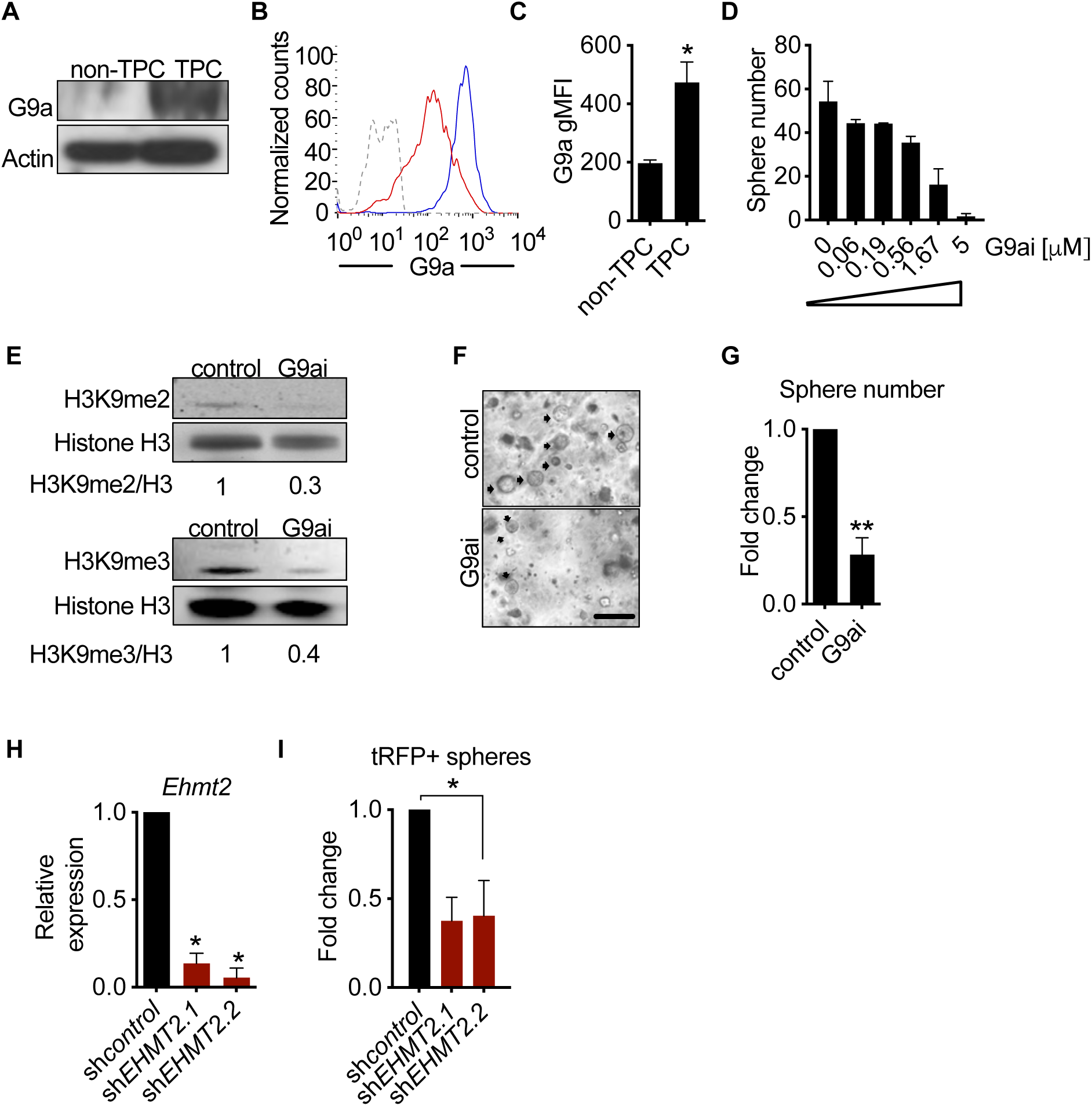
G9a activity is required for *Kras*^*G12D*^;*Trp53* (KP) tumorsphere self-renewal. **(A)** Western blot analysis of G9a in TPC and non-TPC. Actin was used as loading control. **(B)** Flow cytometry analysis of G9a in TPC and non-TPC (blue, TPC, red, non-TPC; grey, Isotype). **(C)** Quantification of G9a geometric fluorescence intensity (gMFI) in (B) (n=2, mean±SEM; two-tailed t-test p=0.05). **(D)** Tumorsphere formation of KP-derived primary cells seeded with increasing doses of G9a inhibitor (n=2, mean±SEM, One-way ANOVA with multiple testing, p<0.005) **(E)** Western blot analysis showing reductions in H3K9me2/3 following G9a inhibitor treatment (G9ai, G9a inhibition) Histone H3 was used as loading control. Ratio of H3K9me to H3 is depicted at the bottom of the western blot. **(F)** Representative image of primary tumorspheres following secondary passaging in the absence of either vehicle control or G9a inhibitor (G9ai, G9a inhibition. Scale bar 100μm). **(G)** Quantitation of tumorsphere growth after secondary passaging (n=5; mean±SEM; two-tailed paired t-test p<0.005). **(H)** Relative qRT-PCR of *Ehmt2* transcripts from primary tumorspheres, expressing either shRNA control (shControl), or shRNAs against *Ehmt2* (shG9a. 1, shG9a. 2), (n=2; mean±SEM; two-tailed paired t-test p<0.05). **(I)** Quantification of turbo RFP (tRFP)-positive tumorspheres following secondary passage of primary tumorspheres expressing control or *Ehmt2* shRNAs. (sh*Ehmt2*.1, n=2; mean±SD, sh*Ehmt2.2*, n=3; mean±SD).

### G9a is required for *in vivo* tumor self-renewal

Previous work has demonstrated that serial re-growth of KP tumors following orthotopic transplantation requires a sustained functional TPC population (Zheng et al., 2013). To evaluate the functional necessity of G9a activity on *in vivo* tumor formation, we evaluated tumor formation following serial transplantation of orthotopically transplanted KP-derived primary cells harboring Doxycycline (Dox)-inducible shRNAs. First, hairpin expression was induced *in vivo* for 13 days by doxycycline administration to mice with established primary lung tumors. Thereafter, shRNA-expressing cells were sorted from primary tumors and assessed for secondary tumorsphere formation *ex vivo* and tumor formation *in vivo* (Figure 2–figure supplement 1A-C). Sorted sh*Ehmt2*-expressing cells from primary recipients showed a significant decrease in both *ex vivo* tumorspheres and *in vivo* secondary tumor formation, establishing a role for G9a in maintaining TPC stemness (Figure 2A-D). Continuous monitoring of secondary transplants *in vivo* revealed a substantial growth impairment in sh*Ehmt2*-expressing tumors, which translated to a significant increase in overall survival (Figure 2E-F). Mice harboring sh*Ehmt2*-expressing tumors eventually succumb to tumor outgrowth; however, analysis of terminal tumors revealed re-expression of *Ehmt2* transcript to a level equivalent to that of control tumors (Figure2–figure supplement 2A,B). This data demonstrates that *Ehmt2* expression is required for TPC-tumor growth. Taken together, these data indicate that G9a activity in TPCs functions to maintain the self-renewal capacity of KP tumors.

**Figure 2.**
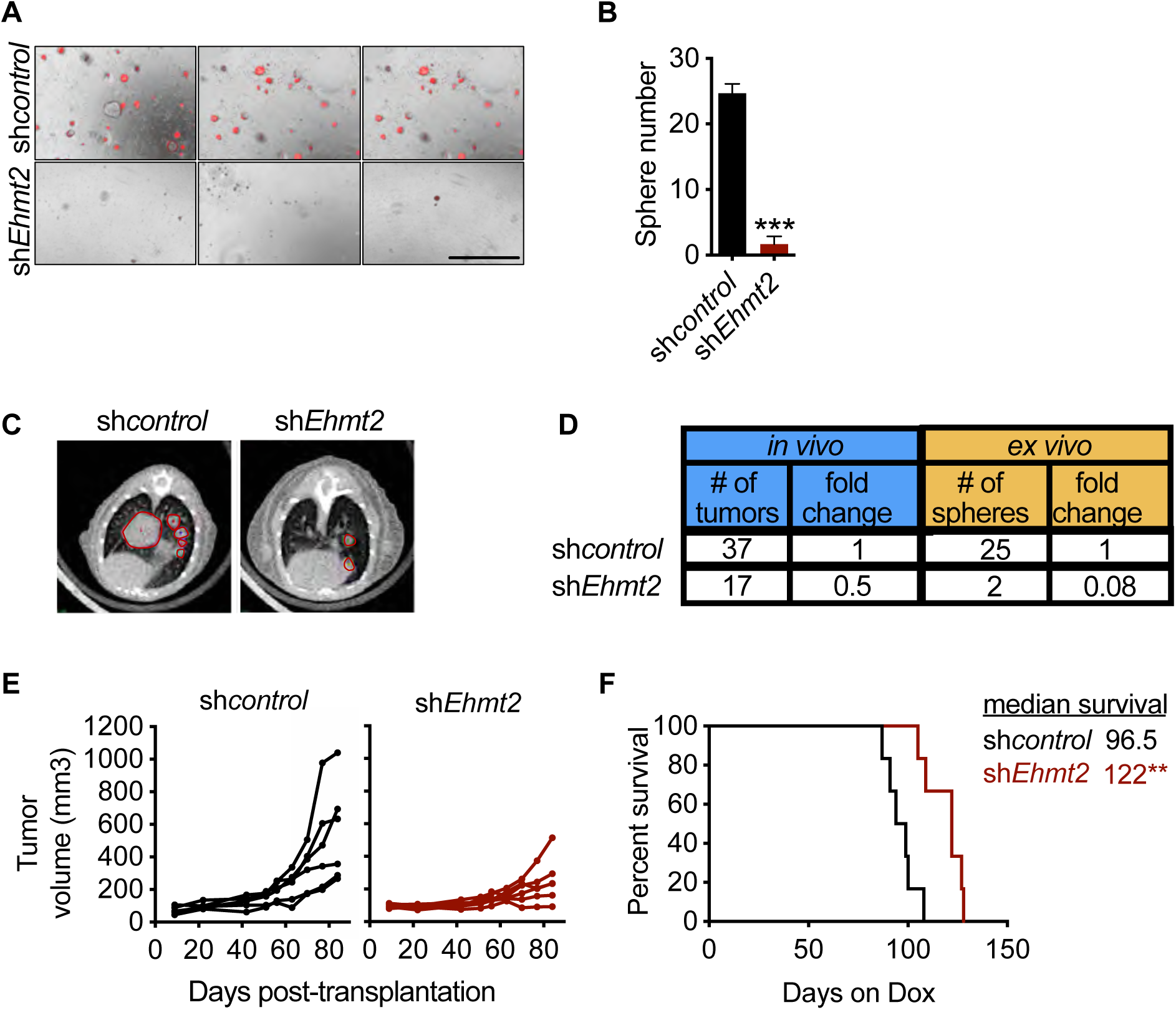
G9a is required for *in vivo* tumor self-renewal. **(A)** *Ex vivo* analysis of tumorsphere formation from primary orthotopic transplanted cells, expressing either shRNA control (sh*control*) or shRNAs against *Ehmt2* (sh*Ehmt2*) (n=3±SD). **(B)** Quantification of tumorsphere formation in panel (A) **(C)** Representative μ-CT images of sh*control*- or sh*Ehmt2*-expressing tumors (n=6) Red circles depicting tumors. **(D)** Table comparing efficiencies of secondary passage *in vivo* and *ex vivo* from orthotopically-transplanted primary KP cells, expressing sh*control* (n=6) or sh*Ehmt2* (n=6). **(E)** Tumor volume in secondary recipient mice orthotopically transplanted with KP cells from primary recipients, expressing either shRNAs targeting control (sh*control*) or *Ehmt2* (sh*Ehmt2.1*) (n=6). **(F)** Graph indicates survival of mice depicted in (E) (n=6 per grouIGeha-Breslow-Wilcoxon test, p<0.005).

### G9a preserves TPC function by preventing terminal AT2 differentiation

To elucidate the mechanistic basis of G9a in maintaining tumor self-renewal, we characterized the phenotypic impact of pharmacological inhibition of G9a in tumorspheres. G9a inhibition resulted in significant reductions in BrdU-labeled cells (5-fold) and expression of cleaved caspase-3 (>10-fold) (Figure3–figure supplement 1A-D). Reduced proliferation and cell death were previously associated with cell differentiation (Domen and Weissman, 1999; Ruijtenberg and van den Heuvel, 2016), therefore, we further explored cell fate changes as a possible treatment outcome. Since lung adenocarcinoma arises predominantly from the distal alveolar compartment (Mainardi et al., 2014; Sutherland et al., 2014; Xu et al., 2012), we quantified established gene signatures pertaining to distal alveolar cell types from RNA sequencing data derived from G9ai-treated tumorspheres (Treutlein et al., 2014). The AT2 gene signature was significantly increased following G9a inhibition, in contrast to other cell lineages (Figure 3A). Protein expression of surfactant protein C (SPC), a canonical AT2 marker was also significantly upregulated in G9a-inhibited tumorspheres (Figures 3A–figure supplement 2), concomitant with an increase in SPC+, we observed an increase in CD74, an additional cell surface marker characterizing AT2 cells (Lee et al., 2013), showing an increase of the double-positive population (1.8-fold) (Figures 3B and 3C). Moreover, we observed a significant increase in the transcript levels of multiple surfactants in both G9a-inhibited and G9a-depleted (sh*EHMT2*) tumorspheres (Figure 3–figure supplement 3A.B), indicating increased/enhanced AT2-like cell fate features when G9a activity is impaired. Importantly, this AT2-like conversion was confirmed in G9a-depleted tumor cells from secondary passage *in vivo* (Figure 3–figure supplement 4). Further evidence supporting cell fate transition was observed when transmission electron microscopy (TEM) of G9a-inhibited tumorspheres revealed a significant increase in lamellar bodies (Balis and Conen, 1964), which are distinct specialized structures responsible for the storage and release of surfactants and serve as a morphometric readout for AT2 cells (Figures 3D and 3E). Together these results demonstrate that G9a inhibition triggers an enhanced/reinforced AT2-like cell state in KP-derived tumorspheres.

**Figure 3.**
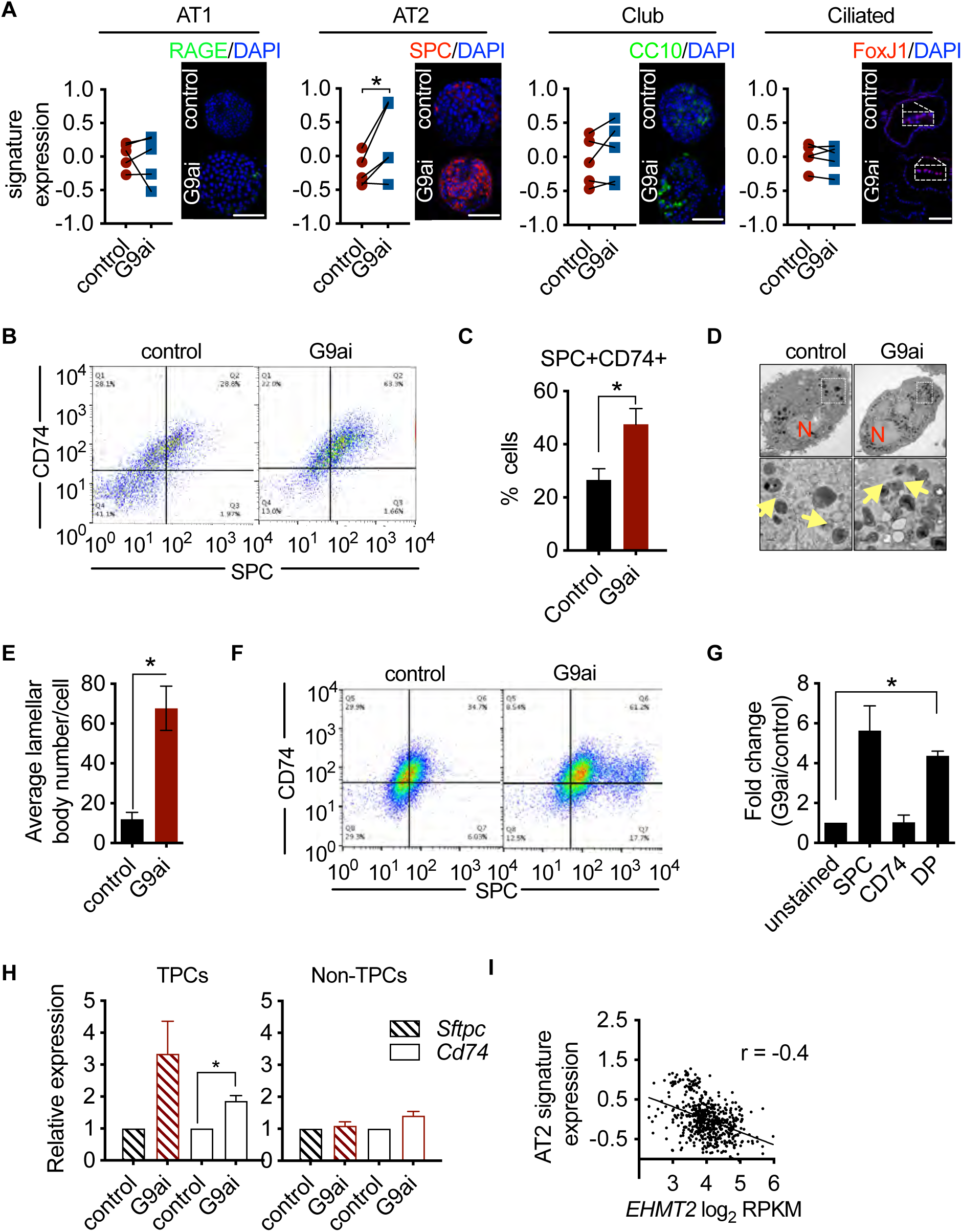
G9a preserves TPC function by preventing terminal AT2 differentiation. **(A)** Graphs show enrichment analyses of distinct alveolar cell-lineage gene signatures in transcriptomes generated from KP-derived primary tumorspheres following G9a inhibition (G9ai) vs. vehicle control (control) (n=4; mean Z-score±SEM, two-tailed paired t-test, p<0.05), each paired with immunofluorescence (IF) micrographs of representative canonical marker from their respective cell lineage. (See Figure (S3E) for quantitation of IF). **(B)** Representative flow cytometry of cells derived from primary tumorspheres treated with either vehicle control (Control) or G9a inhibitor (G9ai) for 5 days and immuno-stained fo23mmuneAT2 markers SPC and CD74. **(C)** Quantification of the SPC-CD74 double positive (DP) population depicted in (B). (n=4; mean±SEM, two-tailed paired t-test, p<0.05). **(D)** Representative transmission electron microscopy (TEM) image of cells extracted from primary tumorspheres, treated as in (B). (Upper panel, scale bar 2μm; lower panel, respective insets in the upper panel, scale bar 0.5μm). (N, nucleus; yellow arrows, lamellar bodies). **(E)** Quantification of TEM in (D) (n=2; mean±SEM, two-tailed paired t-test p<0.05). **(F)** Representative flow cytometry of tumor-propagating cells (TPCs) sorted after G9a inhibitor (G9ai)- or vehicle control-treatment of primary tumorspheres and immuno-stained fo23mmuneAT2 markers SPC and CD74. **(G)** Quantification of (F), showing fold-change in G9ai/control ratio of AT2 markers SPC and CD74 (n=2; mean±SEM, One-way ANOVA with Tukey’s multiple comparison test, p<0.005). **(H)** Relative expression of *Sftpc* and *Cd74* transcripts in G9ai vs. control; TPC and non-TPC respectively. (n=3; mean±SEM, two-tailed paired t-test p<0.05). **(I)** Spearman’s rank correlation analysis between orthogonal human AT2 gene signatures and G9a *EHMT2* transcript in 546 human lung adenocarcinomas. (n=546, linear regression analysis, p<0.0001, r=−0.4)

As G9a activity is critical for the self-renewal of tumorspheres and its expression is selectively enriched in TPCs, we interrogated whether the observed cell fate changes occur within the TPC fraction. G9a inhibition in primary tumorspheres caused a reduction in TPCs, (Figure 3–figure supplement 5), consistent with their reduced stemness. Notably, G9a-inhibited TPCs displayed a statistically significant increase (4-fold) in the AT2 surface markers, SPC and CD74, compared to matched controls (Figures 3F,G). Consistently, *Sftpc* and *Cd74* transcripts increased exclusively in TPCs (Figure 3H). Together, these data indicate that G9a inhibition leads to an induced AT2-like cell state within the TPC subset, thereby reducing their regenerative capacity by a mechanism similar in features to terminal differentiation. To assess whether the relationship between G9a activity and cell state extends to human tumors, we used clinical adenocarcinoma specimens and assessed the association between *EHMT2* transcript and cell lineage gene signatures (Treutlein et al., 2014) in a panel of 546 lung adenocarcinomas (Cancer Genome Atlas Research, 2014). Consistent with our murine data, *EHMT2* expression negatively correlates with the AT2 cell gene signature (Figures 3I–figure supplement 6), supporting the concept that G9a activity impairs terminal differentiation. Taken together, the data indicate that G9a activity represses an alveolar differentiation program in murine lung adenocarcinoma as a means to preserve stem-like properties that enable tumor self-renewal.

### Wnt activation impairs TPC self-renewal and induces AT2 cell lineage marker expression

Previous work indicates that G9a can suppress promiscuous transcription by regulating chromatin structure through the positioning of repressive H3K9 methylation marks in a context-dependent manner (Chen et al., 2012; Kim et al., 2017; Zylicz et al., 2018). We performed an assay for Transposase-Accessible Chromatin using sequencing (ATAC-seq) to characterize chromatin accessibility in G9a inhibitor-treated, tumorsphere-derived TPCs. Surprisingly, very limited changes in chromatin accessibility were observed in TPCs upon G9a inhibition (Figures 4–figure supplement 1A,B), indicating that G9a inhibition does not induce widespread chromatin remodeling: 11 promoter and 320 non-promoter sites became more accessible following G9a inhibition, whereas 3 promoter and 54 non-promoter sites were less accessible (FDR<0.05, fold-change>1.5). The promoter sites were not disproportionally enriched for any pathway, and could not account for the G9a-induced phenotype.

**Figure 4.**
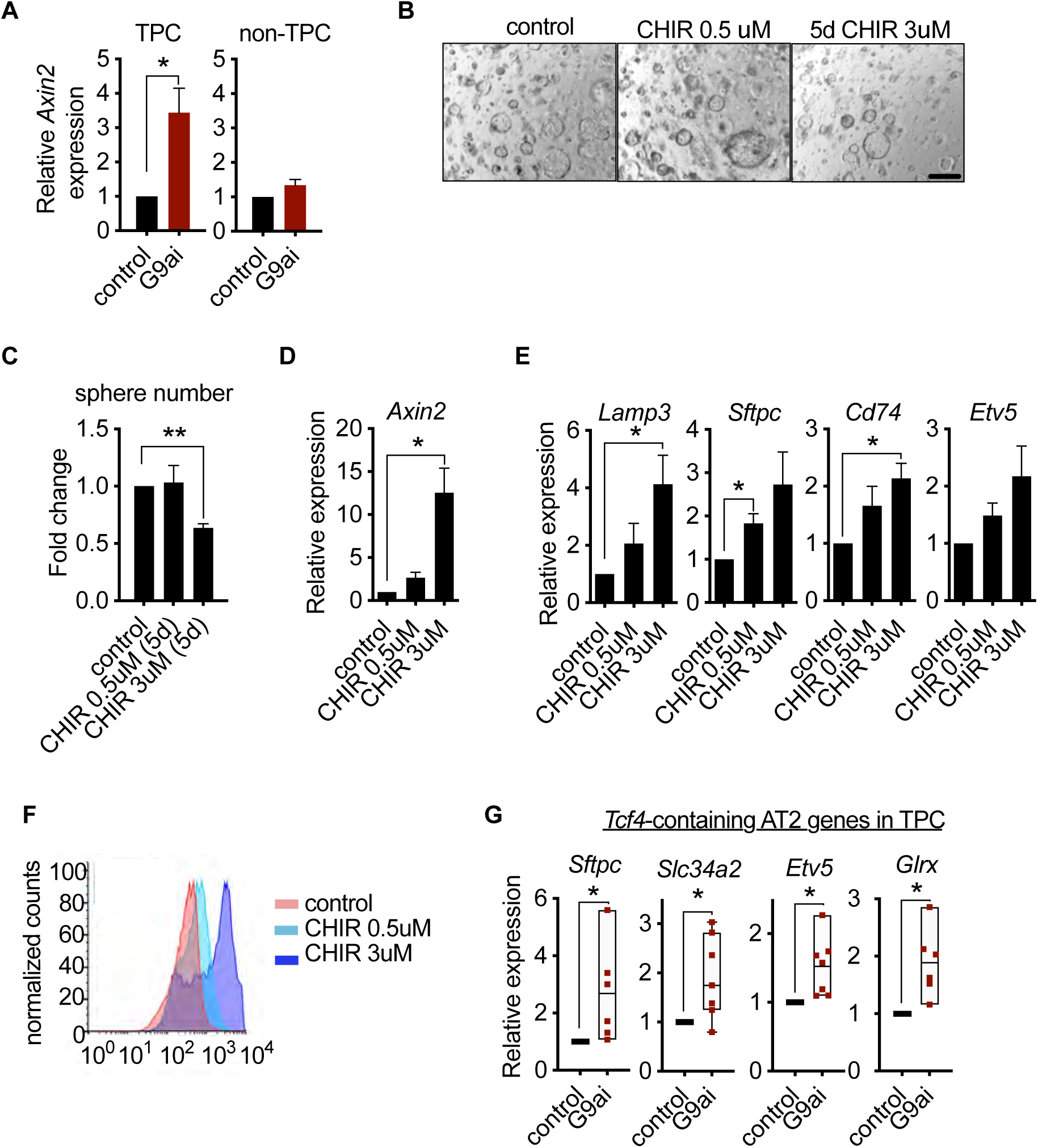
Wnt activation impairs TPC self-renewal and induces AT2 cell lineage marker expression. **(A)** Relative expression of *Axin2* transcripts in tumor-propagating cells (TPCs) and non-TPC following G9a inhibition (G9ai) vs. vehicle control (control) (n=4 mean±SEM; two-tailed paired t-test p<0.05). **(B)** Representative micrographs of primary tumorspheres passaged to single cells following 5 days of the GSK3b inhibitor, CHIR (5d CHIR) at indicated doses vs. vehicle control (control) and assessed for secondary sphere formation (scale bar 100μm). **(C)** Quantification of sphere formation experiments as represented in (b) (n=4 mean±SEM; two-tailed paired t-test p<0.005). **(D)** Relative expression of *Axin2* transcripts in primary tumorspheres (n=5 mean±SEM, two-tailed paired t-test, p<0.05). **(E)** Relative expression of AT2 markers in tumorspheres (n=5 mean±SEM, two-tailed paired t-test p<0.05). **(F)** Flow cytometry for SPC in primary tumorspheres, treated with two doses of GSK3β inhibitor (CHIR) for 5 days vs. vehicle control (control). **(G)** Relative expression of TCF4-containing AT2 markers in TPCs, treated with G9ai vs. control (n=6, mean±SEM, p<0.05).

In the absence of G9ai-induced chromatin accessibility changes, we investigated signaling pathways that influence alveolar cell fate decisions in TPCs. A subset of AT2 cells in normal lung act as tissue stem cells, and Wnt signaling is critical for the maintenance of their stem cell identity (Nabhan et al., 2018; Zacharias et al., 2018). We reasoned that G9a could regulate Wnt signaling as a means to maintain stemness and prevent differentiation in TPCs. Consistent with this line of reasoning, G9a inhibition resulted in a statistically significant increase in *Axin2* expression only within the TPC subset (Figure 4A), implicating a functional link between G9a activity and Wnt signaling. Since G9a inhibition led to *Axin2* upregulation whilst impairing TPC self-renewal and inducing an AT2-like cell fate, we assessed whether Wnt activation alone was sufficient to achieve these outcomes. Indeed, pharmacological activation of Wnt signaling using two doses of the GSK3β inhibitor CHIR99021 (Figure 4B and Figure 4C) impaired TPC stemness as illustrated by reduced tumorsphere self-renewal, consistent with G9a inhibition (Figure 4B-F). Moreover, corresponding Wnt pathway activation and transcriptional activation of AT2 marker genes was observed at secondary passage (Figure 4E), with a concordant dose-dependent increase in SPC surface expression (Fig 4F). To better understand the link between Wnt pathway activation and selective increase of AT2 cell lineage gene expression, we performed an unbiased analysis of transcription factor binding motifs within the promoters of cell lineage signature genes (Treutlein et al., 2014). We found that the *Tcf4* motif ranked 8^th^ among 264 tested motifs for AT2 genes, while ranking much lower for other cell lineages, supporting the concept that AT2 cell fate is directly linked to a Wnt-driven signaling process (Figures 4–figure supplement 2). Moreover, 4 out of 7 *Tcf4*-containing AT2 genes are significantly induced in TPCs upon G9a inhibition (Fig 4G–figure supplement 3), further supporting the concept that the G9a-inhibitor phenotype is Wnt-mediated. Notably, the *Tcf4*-containing AT2 genes that are induced upon G9a inhibition display accessible chromatin configurations at their promoters irrespective of treatment (Fig 4G–figure supplement 4), suggesting that these genes are poised to respond to Wnt-mediated signals and therefore would not require chromatin accessibility changes to enable gene expression (Zacharias et al., 2018).

### G9a restrains Tcf4 gene transcription by repressing chromatin bound β-catenin through RUVBL2

Given the convincing link between inhibition of G9a activity and Wnt-mediated AT2 gene expression, we sought to elucidate the mechanistic basis for Tcf4-mediated gene transcription. Since we observed only a limited change in chromatin accessibility in response to G9a inhibition, we reasoned that a non-histone substrate may be controlling this process. Previous reports have shown that G9a-dependent methylation of the non-histone substrate RUVBL2 (REPTIN, TIP48, TIP49b) can repress HIF1α-mediated transcription (Lee et al., 2010). RUVBL2 has also been shown to antagonize β-catenin activity (Bauer et al., 2000; Mao and Houry, 2017). We reasoned that the non-histone substrate, RUVBL2, might function to repress β-catenin activity through a G9a-mediated mechanism. First, we confirmed the RUVBL2 and β-catenin interaction in a human NSCLC cell line (Figure 5A and 5B). Immunoprecipitation of RUVBL2 showed its ability to interact with β-catenin and HIF1-*α* proteins in hypoxic and normoxic conditions (Figure 5A), consistent with previously reported results (Lee et al., 2010). Reciprocally, immunoprecipitation of β-catenin showed an interaction with RUVBL2. However, the RUVBL2-β-catenin interaction was reduced exclusively in the G9a-inhibited context (Figure 5B), indicating that the RUVBL2-β-catenin interaction requires G9a activity, analogous to that of the RUVBL2-HIF-1*α* (Lee et al., 2010). To better visualize the impact of G9a inhibition on the RUVBL2-β-catenin interaction with the relevant TPC subset, we performed Proximity ligation assay (PLA). Indeed, PLA confirmed the RUVBL2-β-catenin interaction in TPCs and demonstrated a significant loss of signal in the presence of G9ai, providing additional support for the requirement of G9a activity to maintain the RUVBL2-β-catenin interaction (Figure 5C and 5D).

**Figure 5.**
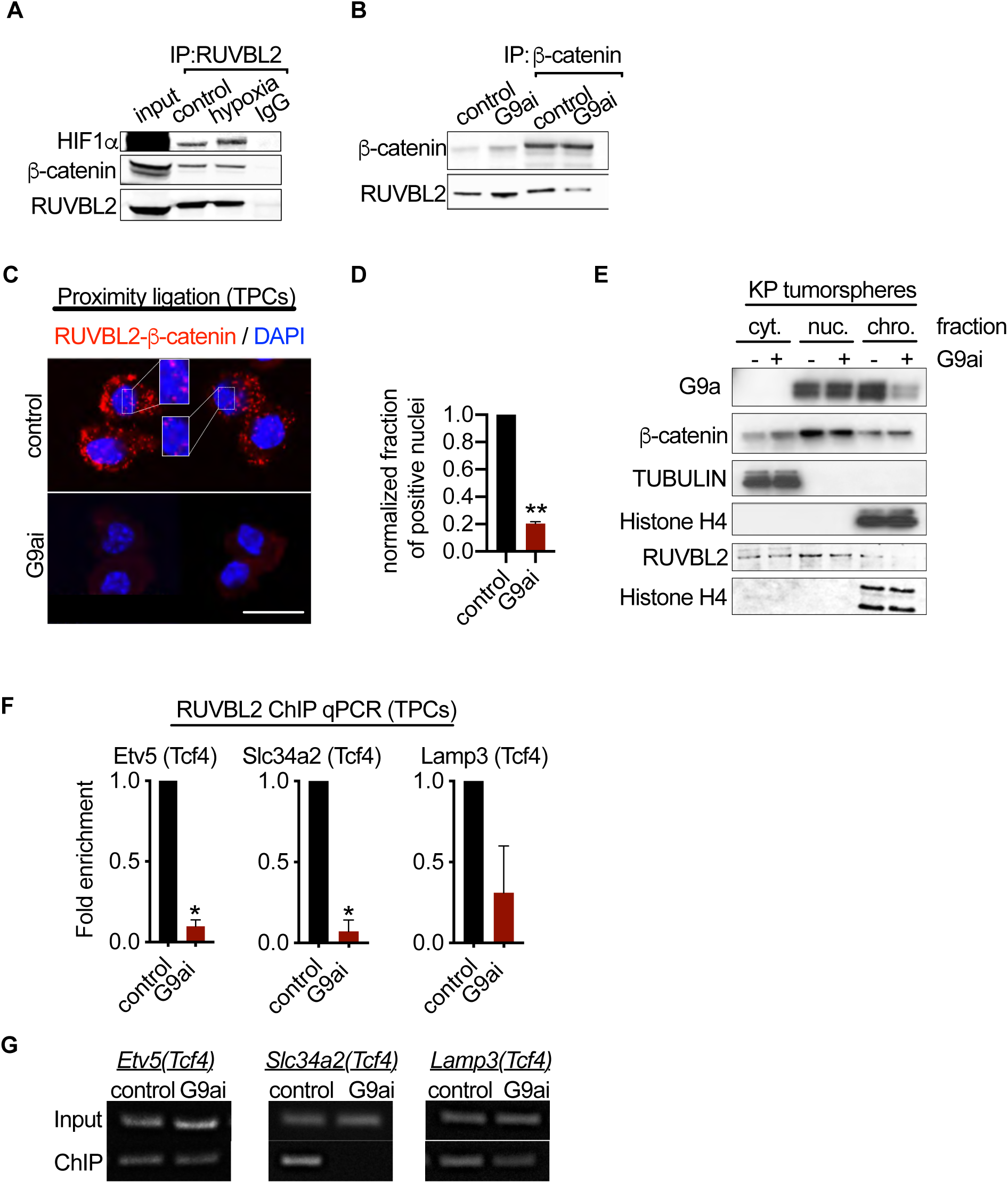
G9a restrains Tcf4-mediated gene transcription by repressing chromatin bound β-catenin through RUVBL2. **(A)** Western blot demonstrating expression of HIF1-*α* and β-catenin in A549 lysates immunoprecipitated with RUVBL2 antibody. Cells were cultured under hypoxic conditions (1% O_2_) vs. control (ambient O_2_). **(B)** Western blot demonstrating co-immunoprecipitation of RUVBL2 in A549 lysates co-immunoprecipitated with β-catenin antibody. Cells were treated with G9a inhibitor vs. control. **(C)** Proximity ligation assay in G9ai-treated vs. vehicle treated TPCs (red, RUVBL2-β-catenin proximity ligation; blue, DAPI counterstain). Insets show a magnification of the red signal in nuclei of vehicle-treated TPCs (scale 10μm). **(D)** Quantification of normalized nuclei with a positive signal inI (n=2; mean±SEM, two-tailed paired t-test p<0.05). **(E)** Cytoplasmic (cyto.), nuclear (nuc.) and chromatin (chrom.) subcellular fractionation of G9ai-treated (as indicated) tumorspheres compared to control. Histone H4 and Tubulin are loading controls of chromatin and cytoplasmic fractions, respectively. **(F)** Chromatin immunoprecipitation using RUVBL2 antibody followed by qPCR (ChIP-qPCR) of areas flanking a *Tcf4* binding sites in promoters of the AT2 genes *Etv5, Slc34a2* and *Lamp3* (n=2 mean±SEM; *Etv5, Slc34a2*, two-tailed paired t-test p<0.05). **(G)** Representative qPCR of *Tcf4* binding motif within promoters of AT2 genes.

In order to gain spatial insight into how G9a inhibition impacts the relationship between G9a, RUVBL2, β-catenin and chromatin, we performed sub-cellular fractionation of tumorspheres. While G9a inhibition showed loss of both G9a and RUVBL2 proteins from the chromatin fraction, equal amounts of β-catenin remain chromatin-bound (Figure 5E). The sustained levels of chromatin-bound β-catenin following G9a inhibition, suggests a critical role for G9a-mediated RUVBL2 regulation that occurs at the chromatin (Figure 5E). Notably, the cytoplasmic fraction of controls confirms the presence of both β-catenin and RUVBL2, consistent with the cytoplasmic signal observed in the TPC PLA control (Figure 5C-E). Given that we established that RUVBL2 interacts with β-catenin and RUVBL2 abundance is reduced in the chromatin fraction when G9a activity is inhibited, we tested whether RUVBL2 chromatin occupancy is changed specifically on *Tcf4*-containing AT2 genes in TPCs. We observed over 90% reduction in RUVBL2 promoter occupancy within *Tcf4* elements of AT2 genes *Slc34a2* and *Etv5* (Figure 5F and 5G). Taken together these results indicate that G9a directly controls *Tcf4*-containing AT2 gene expression through RUVBL2-mediated repression of β-catenin transcription.

### G9a controls Wnt signaling and differentiation within AT2 cells

We next explored whether G9a functions similarly to regulate β-catenin activity in normal AT2 cells. Pharmacological inhibition of G9a *in vivo* for 6 days demonstrated a robust induction of Axin2 protein expression in primary distal alveolar cells using flow cytometry (Figures 6A–figure supplement 1A,B). To assess the impact of G9a inhibitor-mediated Wnt induction on AT2 cell fate, we derived primary AT2 cells from adult murine lung and determined the ability of AT2 progenitors to form alveospheres. *Ex vivo* culturing of primary AT2 cells leads to the formation of alveospheres with cells resembling both AT2 and AT1 cell fates (Barkauskas et al., 2013; Desai et al., 2014). In contrast to KP tumorspheres, inhibition of G9a activity did not impair *ex vivo* alveosphere formation; however, the resulting spheres were significantly smaller relative to controls (Figure 6B). During alveosphere formation and expansion, emerging cells express transcriptional and surface markers consistent with an AT2 to AT1 cell fate transition (Barkauskas et al., 2013; Zacharias et al., 2018). By analyzing expression of surface markers indicative of these fates (Desai et al., 2014), we observed that G9a inhibition significantly reduced the proportion of AT1 cells (SPC-PDPN+) from 67.2% to 24.4% (Figures 6C–figure supplement 2). Interestingly, while the proportion of AT2 cells (SPC+PDPN-) remained unchanged, we observed a marked increase in double positive cells (SPC+PDPN+) from 15% to 62%. Together these data indicate that G9a is required for differentiation of the AT2 progenitor pool and for complete and proper AT1 cell fate transition. Consistent with our previous results in TPCs, G9a inhibition resulted in enhanced Wnt signaling, reflected by increased *Axin2* expression, as well as *Lgr4* and *Lgr5*, two prominent Wnt signaling pathway components (de Lau et al., 2011) (Figure 6D). The observed increase in *Tcf4*-containing AT2 marker gene expression supports increased Wnt-mediated activity in primary AT2 cells as in the TPC context (Figure 6E). These data demonstrate that G9a-mediated regulation of Wnt signaling and cell fate is also observable in untransformed, primary AT2 cells. To test whether G9a loss similarly affects the regenerative capacity of AT2 cells *in vivo*, we deleted G9a in the alveolar compartment by intratracheal administration of an adeno-associated virus encoding *Cre* (AAV9-*Cre*)(Nabhan et al., 2018). This approach allowed us to identify the G9a-targeted cells using a tdTomato reporter and confirm reduced G9a expression from this G9a-deleted population (Figures 6–figure supplement 3). TdT was expressed exclusively in the alveolar space and colocalized preferentially with SPC-expressing cells (Figures 6–figure supplement 4). To further assess how G9a deletion impacts the regenerative capacity of the AT2, we subsequently injured the lung using hyperoxia to promote alveolar repair (Figures 6– figure supplement 5). Four days following injury, G9a loss in the alveolar compartment showed a significant 2.5-fold increase in double positive (SPC+PDPN+) cells, consistent with our ex vivo alveosphere findings (Figure 6F–figure supplement 6). All together these data demonstrate that G9a functions as a cell intrinsic mechanism to directly control cell fate decisions in the context of primary, untransformed AT2 cells.

**Figure 6.**
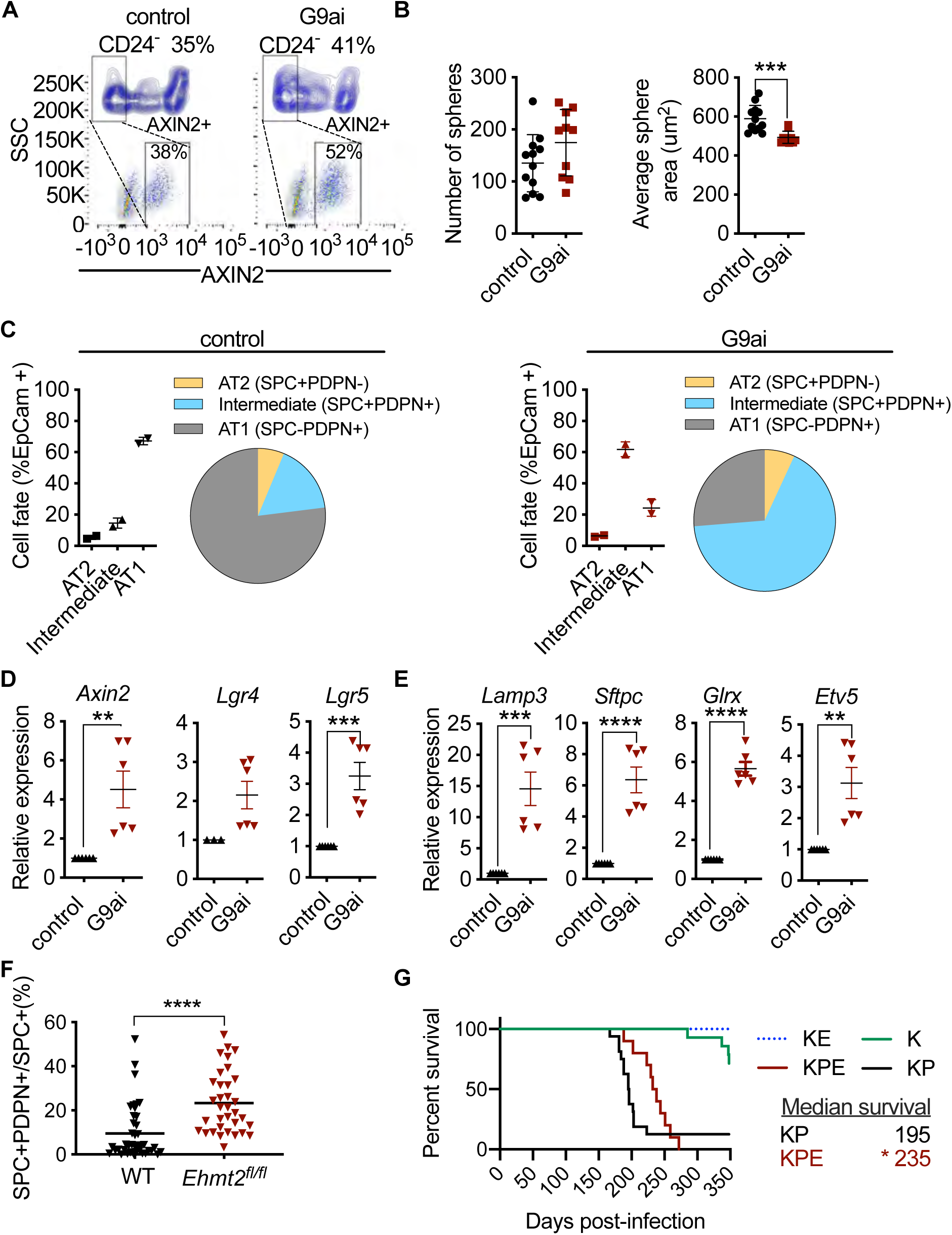
G9a controls Wnt signaling and stemness in AT2 cells and impairs tumor initiation. **(A)** Scatter and contour plots demonstrating intracellular staining of AXIN2 in CD24 negative (CD24^−^) epithelial cells sorted from G9a-inhibited mice compared to control (n=6 for each group). The blue contour plot shows gating on the CD24^−^ cell population. **(B)** Sphere number and size in alveospheres after treatment with G9a inhibitor (G9ai) vs control. (n=10±SEM, two-tailed paired t-test p<0.01). **I** Cell-lineage marker analysis of alveospheres, treated with either vehicle control I or G9ai (D). Each panel shows a graph and a pie-chart depicting epithelial percentages of the AT1 marker podoplanin (PDPN) and the AT2 marker surfactant protein C (SPC). (n=2±SEM, p<0.01). **(D)** Relative expression of *Axin2, Lgr4* and *Lgr5* transcripts (n=6-9±SEM, two-tailed paired t-test p<0.0001). **I** Relative expression of *tcf4*-containing AT2 transcript markers (n=6±SEM; two-tailed paired t-test p<0.05). **(F)** Percentage of SPC+PDPN+ double-positive cells out of SPC+ in WT (n=6) and *Ehmt2*^*fl/fl*^ (n=6) groups, 4 days post hyperoxic (75% O_2_) injury. Quantitation represented as per-field analysis, ≥10,000 cells, scored per field. (n≥10,000, two-tailed paired t-test, p<0.0001 **(G)** Survival of *Kras*^*G12D*^;*Trp53* (KP) (n=16), *Kras*^*G12D*^;*Trp53;Ehmt2*^*fl/f*^ (KPE) (n=10), *Kras*^*G12D*^ (K) (n=14) and *Kras*^*G12D*^;*Ehmt2*^*fl/f*^ (KE) (n=9). Gehan-Breslow-Wilcoxon test, p<0.05).

The impaired regenerative capacity of AT2 cells observed by G9a loss, impelled us to examine the outcome of G9a deletion in Kras-dependent tumor initiation – an event that was previously shown to require cell fate alterations in other tissue contexts (Shibata et al., 2018). To assess the impact on tumor initiation we deleted G9a using a conditional allele of G9a (*Ehmt2* fl/fl) together with conditional *KrasG12D* in the absence or presence of *p53* loss (KPE: *Kras*^*lsl.G12D/wt*^: *Trp53*^*fl/f*l^; *Ehmt2*^*fl/fl*^ and KE: *Kras*^*lsl.G12D/wt*^; *Ehmt2*^*fl/fl*^) KPE mice showed a striking reduction in tumor formation and significant decrease in tumor burden in comparison to KP mice, which translated to a significant increase in overall survival (Figures 6G–figure supplement 7A,B). Of note, the observed increase in overall survival in both KPE and KE mice was independent of p53 status. These results demonstrate that G9a is crucial for Kras-mediated tumor initiation and are consistent with the requirement of G9a to enable AT2 and TPC regenerative capacity and cell fate.

## Discussion

Wnt signals are critical for maintaining or acquiring facultative stem-like properties in lung tissue and distinct disease contexts. Our work reveals a cell-intrinsic mechanism of Wnt pathway activation governed by G9a methyltransferase activity. We find that G9a inhibition directly activates transcriptional activity of chromatin-bound β-catenin within cells, revealing an intracellular mechanism of Wnt signaling control. In the context of lung, G9a activity functions to support regenerative properties of KrasG12D tumors and normal AT2 cells – the predominant cell of origin of this cancer. By discovering this alternative cell-intrinsic mechanism of governing Wnt signaling in stem-like cells, we are able to manipulate cell fate decisions to predictably limit the functionality of these cells to disable tumor propagation

The ability of AT2 stem populations to give rise to differentiated AT2 and AT1 cells is a critical feature that aids in maintaining the integrity of the distal alveolar compartment (Nabhan et al., 2018; Zacharias et al., 2018). How stemness is regulated and maintained is of considerable interest since its dysregulation is an underlying contributor to diseases of the airway, including idiopathic pulmonary fibrosis and cancer. Wnt signaling is a master regulator of AT2 cell fate within the lung, paramount for development, homeostasis and damage repair. Regulation and tuning of Wnt signals are necessary to ensure proper repair in the alveolar compartment, but the mechanisms are context dependent. For example, β-catenin loss in AT2 cells enhances AT1 conversion, indicating Wnt signaling functionally prevents differentiation of alveolar AT2 stem cells (Frank et al., 2016; Nabhan et al., 2018) Our findings are consistent with this paradigm as functional perturbation of G9a activity enhances Wnt signaling, indicating impairment of AT2 plasticity, as well as the regenerative capacity of TPCs. Recent reports (Nabhan et al., 2018; Zacharias et al., 2018; Zepp et al., 2017) further describe how Wnt regulation is remarkably nuanced within the distal lung with the discovery and characterization of distinct niches that provide exogenous Wnt ligands to support neighboring stem cells. In the context of tissue damage, these interactions are likely disrupted and thereafter differentiated cells can adopt a facultative state defined by changes in Wnt signaling. Although autocrine Wnt secretion has been proposed as a mechanism to self-sustain the Wnt signaling requirements while outside of the niche, our discovery provides a means to enable cell intrinsic regulation of β-catenin transcription, providing a safe switch to maintain AT2 cell identity when a mesenchymal Wnt-providing niche, is no longer tethered to the epithelial cell.

In lung tumorigenesis, we observe that G9a loss at the time of tumor initiation leads to a significant reduction in tumor formation, as well as, tumor burden resulting in significantly increased overall survival. Of note, genetic cooperativity between Ras and Wnt signaling pathways has been reported (Pacheco-Pinedo et al., 2011) and linked to cell fate effects. In the context of *Kras*-mutant lung tumor initiation, genetic activation of Wnt signaling using mutant beta catenin within *Scgb1A1+* cells leads to a distal cell fate change consistent with our results. However in contrast to our AT2-derived tumor-initiation model, results in enhanced tumor formation within *Scgb1A1+* expressing cells (Pacheco-Pinedo et al., 2011). Notably, this work differs in the cell of origin, but in that regard, it is not yet clear how distinct, alternate thresholds of Wnt signaling contribute to determining cell fate within different cell lineages in the lung. Moreover, Pacheco-Pinedo et al., used a stable form of beta catenin, whereas loss of G9a activity leads to signaling of chromatin-bound beta catenin, which likely differ in both signal strength and duration. Others have reported the necessity of Wnt signaling to maintain tumors and suggested that Wnt gradients might be critical for self-renewal, in line with the necessity for a specific Wnt threshold to support this process (Tammela et al., 2017). Of note, a recent report surprisingly demonstrated enhanced tumor formation when initiating tumors with mutant Kras and G9a knockdown (Rowbotham et al. 2018), This is in direct contradiction to our findings using a genetic model to delete G9a concurrent with *Kras*^*G12D*^ in tumor initiation. Importantly, the shRNA seed sequence used in the Rowbotham study targets nine other target transcripts with 100% homology *in addition to Ehmt2* (Table S1), raising the possibility that genes other than *Ehmt2* may be implicated in the described phenotype. Notably, one of the genes targeted by this shRNA construct is *BABAM1*, which is known to increase the metastatic capability of a murine Kras-mutant lung cancer-derived cell line (Chen *et al.*, Cell 2015; PMID: 25748654, (Table S1). While in contrast to our findings, these technical differences preclude definitive conclusions from those experiments within their study.

Our work implicates G9a as an important mediator of Wnt signals in order to maintain tumor propagating capacity. This cell autonomous mechanism for activating Wnt signaling may be beneficial in contexts of tumor seeding and regrowth, allowing independence from niche factors. G9a activity functions similarly in primary AT2, as well as, *Kras*-transformed tumor cells to maintain stemness in both contexts, suggesting that TPCs may share functional features of their respective cell of origin. The discovery that G9a functions to directly regulate Wnt signaling deepens our mechanistic understanding of G9a activity and presents potential opportunities for targeting in both lung cancer, as well as other AT2-mediated lung pathologies.

## Materials and Methods

### Mice

*Kras*^*LSL-G12D*^ (Jackson et al., 2001), *Trp53*^*flox/flox*^ (Jonkers et al., 2001), *Trp53*^*frt/frt*^ (Lee et al., 2012), *Rosa26*^*LSL-tdTomato*^ (Madisen et al., 2010) were licensed by Genentech Inc. All animal studies were approved by the Institutional Animal Care and Use Committee at Genentech and adhere to the Guidelines for the Care and Use of Laboratory Animals. Tumors were induced in KP mice at 8–12 weeks of age using intranasal infection of AdCMV-Cre or Adeno-Flp-IRES-Cre (Baylor College of Medicine) at 1.2 × 10^7^ plaque-forming units (PFU).

### Orthotopic transplantation studies

For tumor formation, 8-12 week-old recipient mice were intratracheally transplanted with 25,000-45,000 inducible shRNA-carrying primary KP tumor cells per mouse. Cells were resuspended in 60µL MEM alpha (Gibco) before transplantation and monitored using micro CT as previously described(Zheng et al., 2013).

### Cell isolation and tumorsphere preparation

Pooled KP tumors from 3-6 mice per experiment, were extracted out of the lung and completely minced with a razor blade. The material was resuspended in DMEM-F12 media containing 10% FBS, 1X P/S and 2µg/ml Collagenase-Dispase in 50 ml conical tubes and incubated for 1-1.5h in a 37°C incubator on a shaking platform. Digested material was sequentially filtered through 70 micron and 40 micron strainers, distributed into 15 ml conical tubes and centrifuged for 5 min at 500g. Pellets were resuspended in hypotonic lysis buffer (15mM NH_4_Cl, 10mM KHCO_3_, and 0.1mM EDTA) for 1-2 minutes, neutralized with DMEM-F12 and spun down on a 1ml FBS cushion to remove cell debris. Final cell pellets were resuspended in PBS containing 10% FBS. For tumorsphere assays, KP tumor cells were mixed with Matrigel in tumorsphere media as previously described(Zheng et al., 2013). TPCs were sorted and analyzed using the influx machine (BD). Sorting was performed as previously described by (Zheng et al., 2013) with the following modifications. Immune cell lineage content was excluded by sorting with the following <u>antibodies:</u> biotin-conjugated CD45 (BD, 553078, 30-F11,1:200), CD31(BD, 553371, MEC13.3, 1:200), Ter119 (BD, 553672, Ter119, 1:200), thereafter Biotin-conjugated antibodies were detected with Phycoerythrin (PE)/Cy7 Streptavidin (Biolegend, 405206: 1:300), and subsequently stained for EpCAM+ FITC (Biolegend, 118208, G8.8, 1:20) and TPC markers CD24 PerCP-eFluor 710 (eBioscience, 46-0242, M1/69: 1:300), ITGB4-PE (Biolegend, 123602, 346-11A, 1:20) and Notch1 (Biolegend, 130613, HMN1-12, 1:80), Notch2 (Biolegend, 130714, HMN-2-35, 1:80), Notch3 (eBioescience, 17-5763-82, HMN3-133, 1:80), Notch4 (Biolegend, 128413, HMN4-14, 1:80). All anti Notch antibodies are Allophycocyanin (APC) conjugated and used as a pool.

### TPC gating strategy

Dead cells and doublet cells were excluded from the sort respectively. EpCAM-Lineage scatter plot was used to gate on epithelial cells. Each TPC marker was gated in an individual scatter plot against a mock gate in the following sub-gating scheme: CD24+ cells gated from EpCAM+ cells, ITGB4+ cells gated from CD24+ cells and Notch^HI^ gated from ITGB4+ cells. Non-TPCs were classified as the ‘non-positive’ cells for each individual marker scheme and gated accordingly. All non-TPC were combined in one single tube.

### Tumorsphere and TPC assays

For G9a inhibition assays in KP tumorspheres, 10,000-20,000 KP primary tumor cells from 3-6 mice for each biological replicate were seeded in Matrigel for 4-5 days before treatment. Each biological replicate contained n=3-4 technical replicates. Tumorspheres were then treated with either vehicle or 2 µM of the G9a inhibitor Unc0642 (Bio-techne) for an additional 5-7 days. For Wnt activation assays, established tumorspheres were treated with either 0.5 µM or 3 µM CHIR99021 (Tocris) for 5 days. For G9a inhibition assays in TPCs, 5 ×10^6^-10 ×10^6^ KP primary tumor cells from 6-8 mice per experiment were seeded per 5 ml Matrigel plug for 4-5 days before treatment and treated as described above. Cells were then extracted out from Matrigel as single cell suspensions and sorted for the TPC cell subset. TPC isolation directly from treated or untreated cultured tumorspheres was performed using the Influx sorter (BD).

### Secondary sphere formation assays

For secondary sphere formation assays following G9a inhibition or depletion, established KP tumorspheres were mechanically and enzymatically dissociated with 2µg/ml collagenase/Dispase (Roche). To obtain single-cell suspensions, dissociated KP tumorspheres were further treated with Accutase (Corning) for 5-10 min, counted and re-seeded in Matrigel in 3-4 technical replicates per experiment. Matrigel, spheres were then counted using a 10×20 eyepiece containing 0.5×0.5mm grid.

### Alveolar and AT2-derived alveosphere preparations

AT2 derived alveosphere assays were generated by isolating Epcam^+^/CD24^−^ cells as previously described(Barkauskas et al., 2013; McQualter et al., 2010). Sorted cells were either intracellularly stained for Axin2 Abcam (109307, EPR2005, 1:100) or alternatively, resuspended in SAGM media (Lonza) and mixed in a 1:1 ratio with growth factor-reduced Matrigel (BD Biosciences). 100 μl of mixed cells/Matrigel suspension was placed in 24-well Transwell inserts (Falcon) and allowed to solidify. To allow alveospheres growth, 5 ×10^4^ MRC5 human lung fibroblasts (ATCC CCL-171) were seeded in the bottom chamber supplied with 500 μl MTEC media. Media was changed every other day. G9a inhibitor was replaced every 2-3 days.

### IP and ChIP (chromatin immunoprecipitation)

For Co-IP experiments β-catenin and RUVBL2 were pulled-down from vehicle or G9a inhibitor-treated A549, using Thermo Fisher (71-2700) or Bethyl (A5302-537A) antibodies respectively, and utilizing the Dynabeads Co-IP kit (14-321) as indicated by protocol. To detect interacting proteins, membrane was blotted for β-catenin BD (610153) or RUVBL2 Bethyl (A302-536A). ChIP experiments were performed on tumor propagating cells (TPCs) sorted directly from vehicle or G9a inhibitor treated-tumorspheres using the Diagenode True-ChIP protocol (C01010140) with the following modifications. Briefly, 50K-100K cells were crosslinked with 1% formaldehyde for 8 min, chromatin was sheared to 100-500bp fragments using the Bioruptor Pico sonicator for 3 cycles of 30 sec on/off in 0.5ml tubes (Diagenode). Quantitation of chromatin was performed using a qubit fluorimeter where at least 10 ng of chromatin were used per experiment. Pulldown was performed with 0.5ug RUVBL2 Antibody Bethyl (A302-537A) overnight then incubated with 30 min pre-blocked protein A conjugated Dynabeads. Immunoprecipitated material was then washed as indicated by protocol with an additional LiCl buffer wash (0.25M LiCl, 1% IGEPAL, 1% deoxycholic acid, 1mM EDTA, 10mM Tris, pH 8.1). Samples corss-links where then reversed and then samples were pre-amplified using PerfeCTa, QuantaBio (95146-005). The precipitated DNA and input DNA were quantified by qPCR using specific primers flanking a TCF4 binding site in the promoters of the following genes, *Etv5, Slc34a2, Lamp3.*

### Cell lineage marker analysis

Mouse cell lineage markers were obtained from Treutlein et al., 2014(Treutlein et al., 2014) using the following selection criteria: Pearson correlation >=0.5, p-value (GBA, BH corrected) <0.05. This resulted in 30 mouse genes characteristic of AT1 cells, 28 genes for AT2 cells, 43 genes for Club cells, and 84 genes for Ciliated cells. The biomaRt R package was used to map mouse genes to human orthologs, resulting in 28 human AT1 markers, 26 AT2 markers, 39 Club markers, and 65 Ciliated markers. For the assessment of lineage markers in human and mouse samples, expression data Z scores were first calculated for each gene across the 546 human lung adenocarcinoma tumors, and across the 10 mouse KP tumorsphere samples. Cell lineage signature scores were then calculated for human adenocarcinoma tumors as the average z-score expression of the human markers for each cell lineage and for mouse samples as the weighted average z-scored expression of the mouse markers with weights set to −log10(p-value). P-values were obtained from Table S4(Treutlein et al., 2014). **Assay for Transposase-Accessible Chromatin using Sequencing (ATAC-Seq)** Cells were aliquoted into cryovials, frozen, and shipped to Epinomics, Menlo Park, CA. Cells were processed as previously described(Buenrostro et al., 2013). Paired-end reads were aligned to mouse reference genome GRCm38 using GSNAP(Wu and Nacu, 2010) version ‘2013-10-10’, allowing a maximum of two mismatches per read sequence (parameters: ‘-M 2 -n 10 -B 2 -I 1 –pairmax-dna=1000 –terminal-threshold=1000 –gmap-mode=none –clip-overlap’). Duplicate reads and reads aligning to locations in the mouse genome containing substantial sequence homology to the mitochondrial chromosome or to the ENCODE consortium blacklisted regions were omitted from downstream analyses. Remaining aligned reads were used to quantify chromatin accessibility according to the ENCODE pipeline standards with minor modifications as follows. Accessible genomic locations were identified by calling peaks with MACS 2.1.0(Zhang et al., 2008) using insertion-centered pseudo-fragments (73bp – community standard) generated on the basis of the start positions of the mapped reads and a width of 250bp. Accessible peak locations were identified as described; briefly, we called peak significance (cutoff of p=1e-7) on a condition-level pooled sample containing all pseudo-fragments observed in all replicates within each condition. Peaks in the pooled sample, independently identified as significant (cutoff p=1e-5) in two or more of the constituent biological replicates were retained, using the union of all condition-level reproducible peaks to form the atlas. (https://www.encodeproject.org/atac-seq/). The atlas consisted of 184,032 peaks, of which 37,551 located in promoter regions.

Chromatin accessibility within each peak for each replicate was quantified as the number of pseudo-fragments overlapping a peak and normalized these estimates using the TMM method(Robinson and Oshlack, 2010). Differentially accessible peaks between control and G9a-inhibited samples were identified using the framework of a linear model, accounting for TPC and non-TPC and implemented with the edgeR R package. Significant differences in chromatin accessibility levels within a peak between G9ai-treated and control-treated samples was set to log2-fold change > 1.5 and FDR < 0.05. The fold-change of ATAC-seq peaks were used as input for gene set enrichment analysis using the Hallmark MsigDB gene set collection, v6.1(Liberzon et al., 2015).The Integrative Genomics Viewer (IGV) was used for visualization of ATAC-seq peaks near genes of interest(Robinson et al., 2011). All source code and sequencing data are available at https://github.com/anneleendaemen/G9a.CellIdentity.Lung

## Acknowledgments

We appreciate the support of all the Junttila lab members. We would like to thank Elaine Storm and the De Sauvage lab for providing critical reagents. We are grateful to the sequencing lab for their superb work. We also received technical support from the in-house genotyping and murine reproductive technology core groups, especially Tiffany Yuan. We would like to acknowledge the laboratory animal research core for facilitating the support needed to enable *in vivo* experimentation, as well as, the center for advanced light microscopy for providing help with immunofluorescence quantitation and all FACS lab members, especially Jonathan Paw and George Tweet who were instrumental with devising critical sorting strategies.

## Author Contributions

A.P and M.R.J. wrote the manuscript. A.P. and M.R.J designed experimentation. A.P. performed all tumorsphere experimentation and analysis. T.L. and A.P. performed *in vivo* experimentation. X.W. performed alveosphere experimentation. A.D., and M.H., contributed computational analysis. C.P. helped with FACS sorting. Z.M. provided support for both RNA and ATAC sequencing. A.K.M., performed transmission electron microscopy scanning and analysis. O.F. provided pathology support. J.E. quantitated the immunofluorescent images. B.H. supported construct design. J.T.G. assisted with ATAC-seq analysis. E.L.J and M.R.J supervised the project.

## Supplementary Figures

**Figure 1–figure supplement 1.**
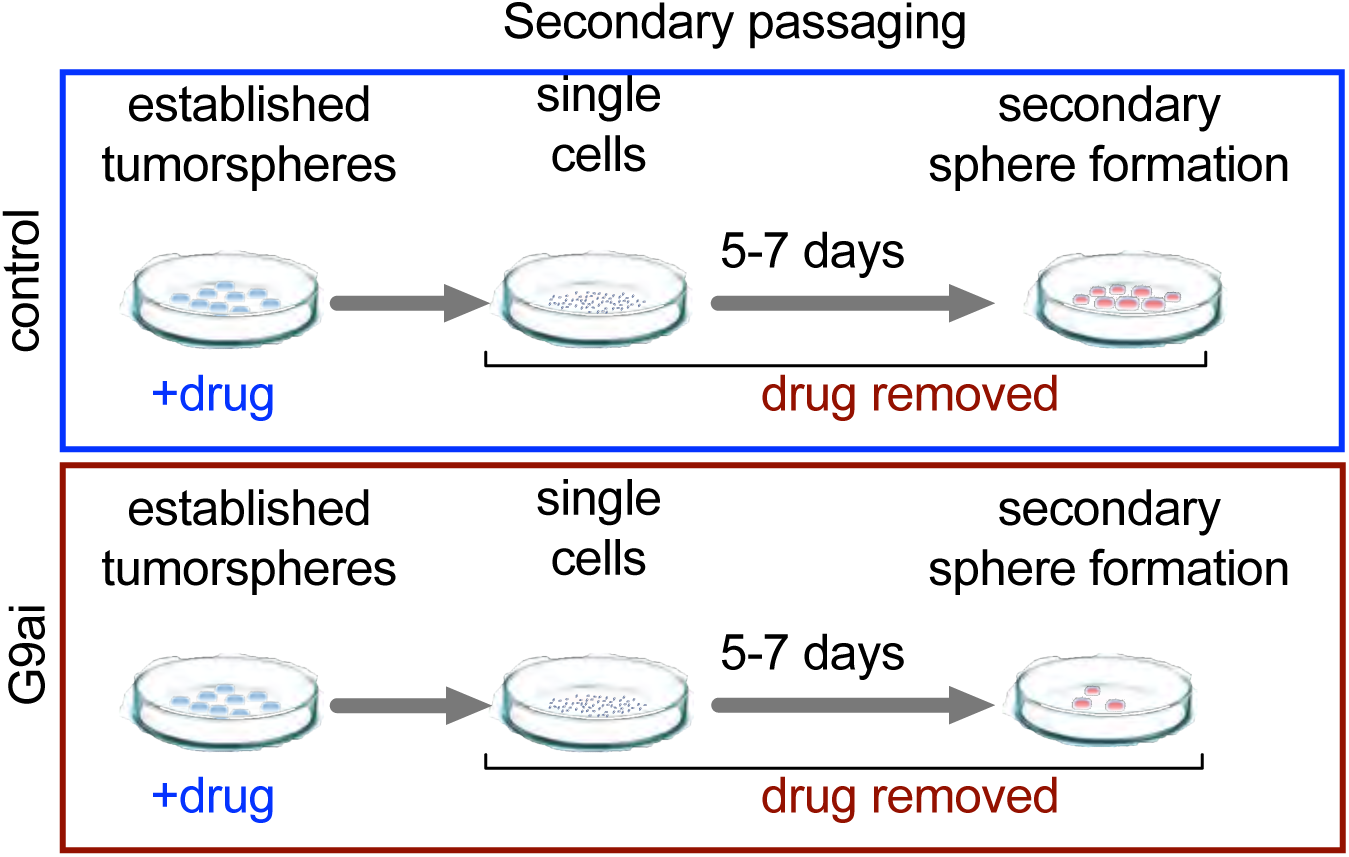

**Figure 2–figure supplement 1.**
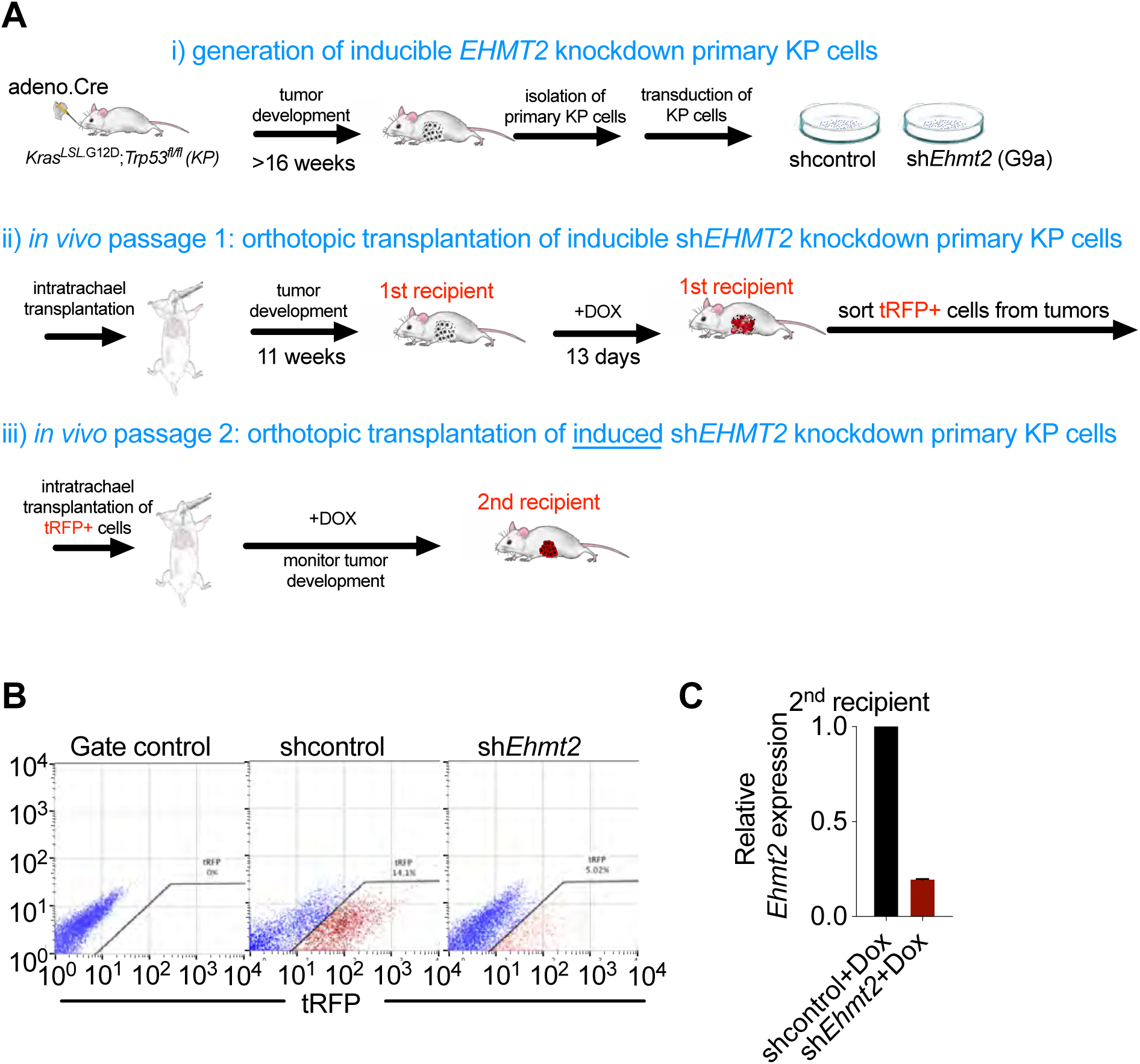
Analysis of *in vivo* serial orthotopic transplantation of primary KP cells harboring *EHMT2* targeting hairpins. **(A)** Schematic representation of serial orthotopic transplantation of primary KP cells can be viewed in three basic parts: i) Primary tumors are initated via intranasal infection of adenovirus expressing Cre recombinase in Kras^LSL.G12D^; p53^flox/flox^ (KP) mice. Tumors develop with a latency of approximately 16 weeks. Primary tumors are isolated from KP mice and immediately transduced with lentiviral constructs harboring doxycycline-inducible hairpins and a tRFP label to facilitate identification of hairpin-expressing cells. Ii) Transduced primary KP cells are then orthotopically seeded into the lungs of wild-type recipient mice via intratracheal administration. Animals are thereafter monitored for tumor formation using microCT. Once tumor formation is confirmed in 1^st^ recipient mice, animals are stratified and dosed with doxycycline for 13 consecutive days to induce expression of latent hairpins targeting either *EHMT2* or control transcripts. After 13 days of expression, tumors from 1^st^ recipients are harvested and tRFP+ cells are sorted to identify hairpin expressing cells. Iii) tRFP+ cells are then orthotopically transplanted into a 2^nd^ recipient. The animals are maintained on Doxycycline and tumor growth is monitored. **(B)** Flow cytometry showing gating strategy of primary tRFP-sorted cells prior to secondary transplantation (left plot, tRFP gate control; middle and right plot, sorted tRFP-positive cells from shcontrol and sh*EHMT2*, respectively). **(C)** Relative expression of sh*Ehmt2.2* in tRFP-sorted tumor cells prior to secondary transplantation (n=6).

**Figure 2–figure supplement 2.**
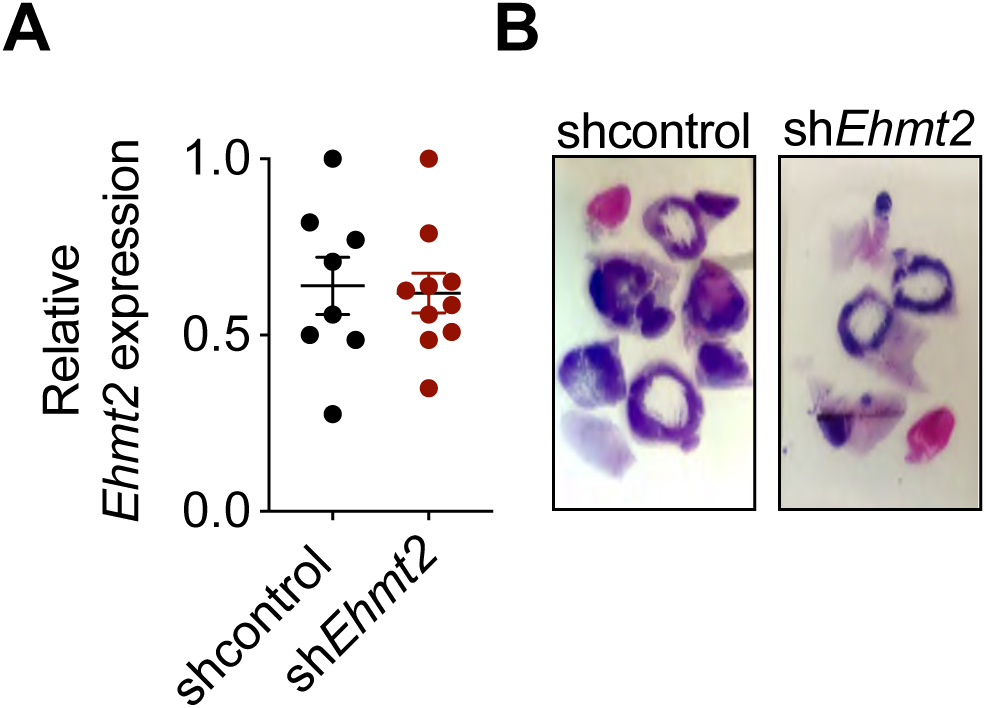
Analysis of terminal tumors from *in vivo* serial orthotopic transplantation of primary KP cells harboring *EHMT2* targeting hairpins. **(A)** Relative expression of *Ehmt2* transcripts from tumors of shcontrol and sh*EHMT2.2* secondary recipients at study termination, showing no statistical difference (Each point represents one tumor from n=9 for each group). **(B)** Micrographs showing extracted tumor areas taken for transcript expression analysis from both control and *shEhmt2* secondary recipient tumor transplants.

**Figure 3–figure supplement 1.**
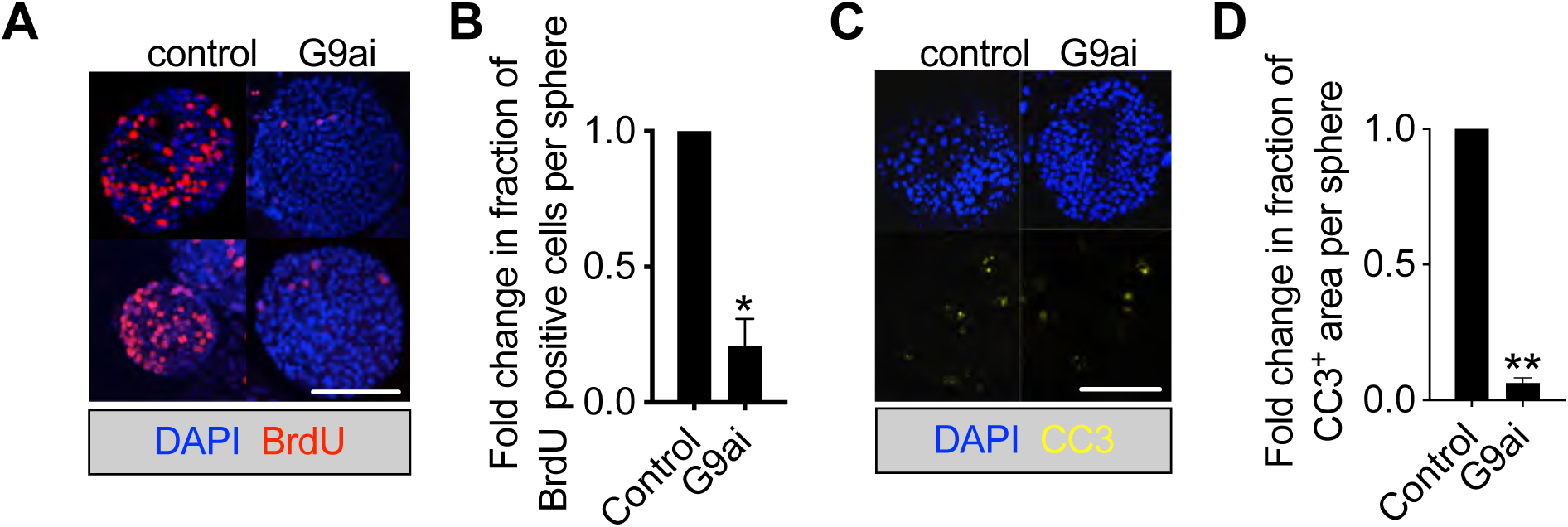
G9a inhibition reduces apoptosis and proliferation in tumorspheres. **(A)** Proliferation in KP-derived primary tumorspheres demonstrated by BrdU staining, following 5 days with G9a inhibitor (G9ai) or vehicle control (control) (red, BrdU immunofluorescence; blue, DAPI counterstain; scale 500μm). **(B)** Quantification of BrdU-positive nuclei depicted in (a) (two-tailed paired t-test; n=3; mean±SEM, p<0.05). **(C)** Micrographs indicate apoptosis (CC3, Cleaved caspase-3) in primary tumorspheres, following 5 days with G9ai or control (blue, DAPI counterstain; scale bar 500μm). **(D)** Quantification of CC3-positive area per sphere area, depicted in (c). (two-tailed paired t-test n=2; mean±SEM, p<0.01).

**Figure 3–figure supplement 2.**
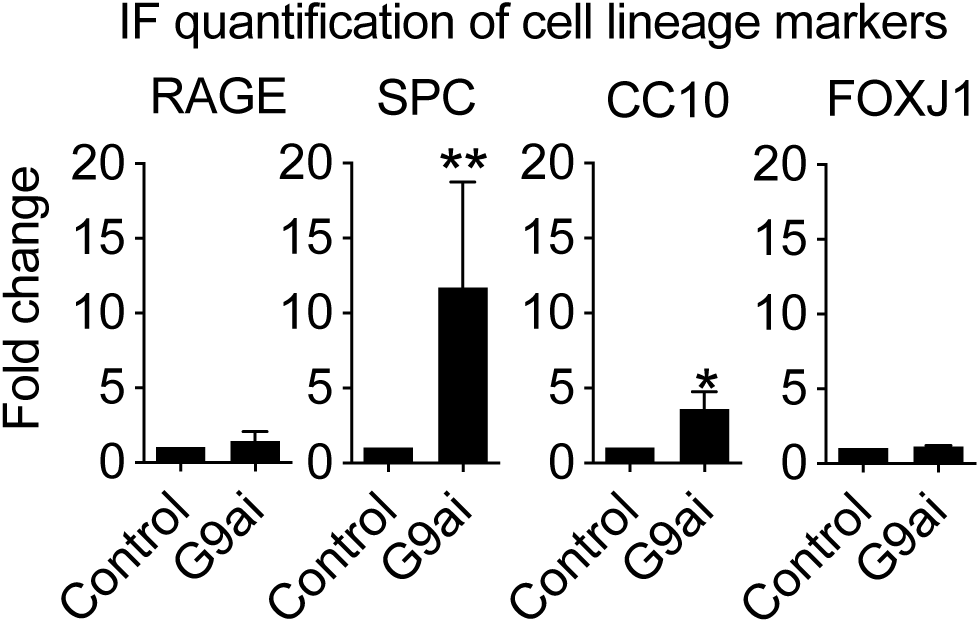
Quantification of immunostaining in control and G9ai-treated tumorspheres (RAGE: control, n=28; G9ai, n=21. SPC: control, n=15; G9ai, n=9; two-tailed paired t-test, p<0.005. CC10: control, n=15; G9ai, n=9; two-tailed paired t-test, p<0.05. FoxJ1: control, n=28; G9ai, n=21).

**Figure 3–figure supplement 3.**
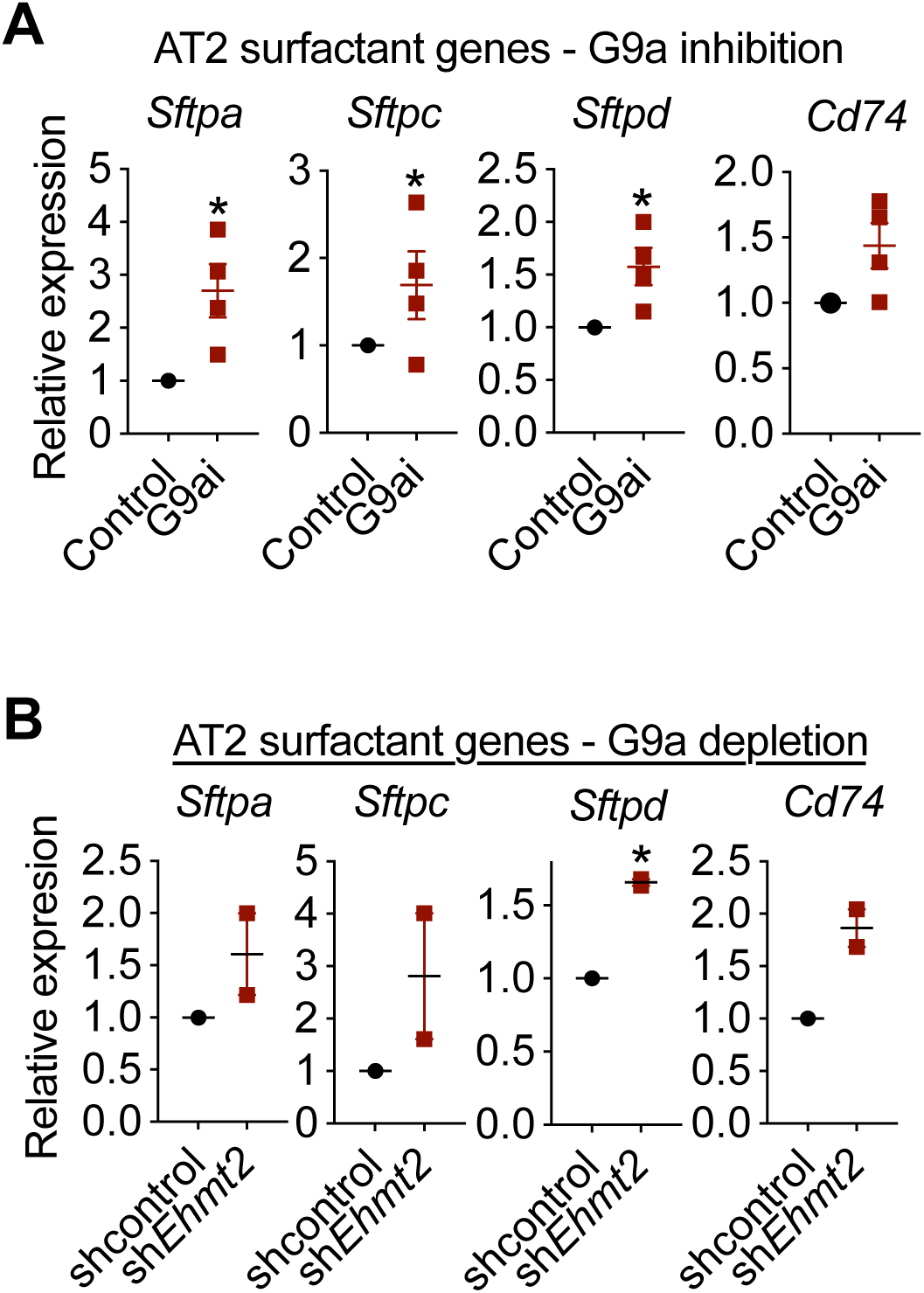
Induction of surfactants following G9a depletion and pharmacologic inhibition. **(A)** Relative expression of surfactants and *Cd74* transcripts following G9a inhibition (n=4±SEM; p<0.05) or sh*Ehmt2* vs. shcontrol (control) (two-tailed paired t-test n=4±SEM; p<0.05) **(B)** Relative expression of surfactants and *Cd74* transcripts following G9a inhibition (n=4±SEM; p<0.05) or sh*Ehmt2* vs. shcontrol (control) (n=2±SEM)

**Figure 3–figure supplement 4.**
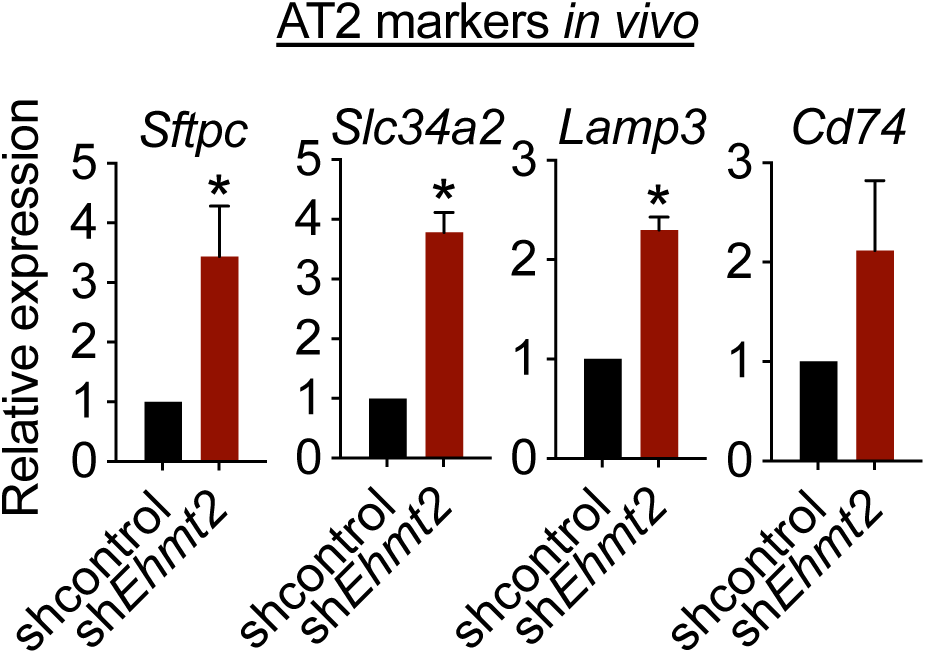
Relative expression of AT2 markers in tRFP-sorted tumor cells derived from primary recipients expressing either shcontrol or sh*Ehmt2* (n=3; mean±SEM;,*Sftpc, Slc34a2*, Lamp3, Cd74; two-tailed paired t-test p<0.05).

**Figure 3–figure supplement 5.**
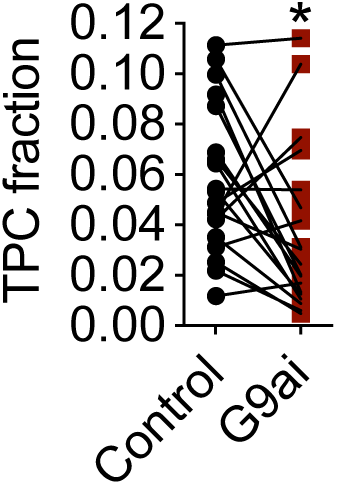
Quantification of TPC fraction following G9ai vs. control (n=19; mean±SEM; two-tailed paired t-test p<0.05).

**Figure 3–figure supplement 6.**
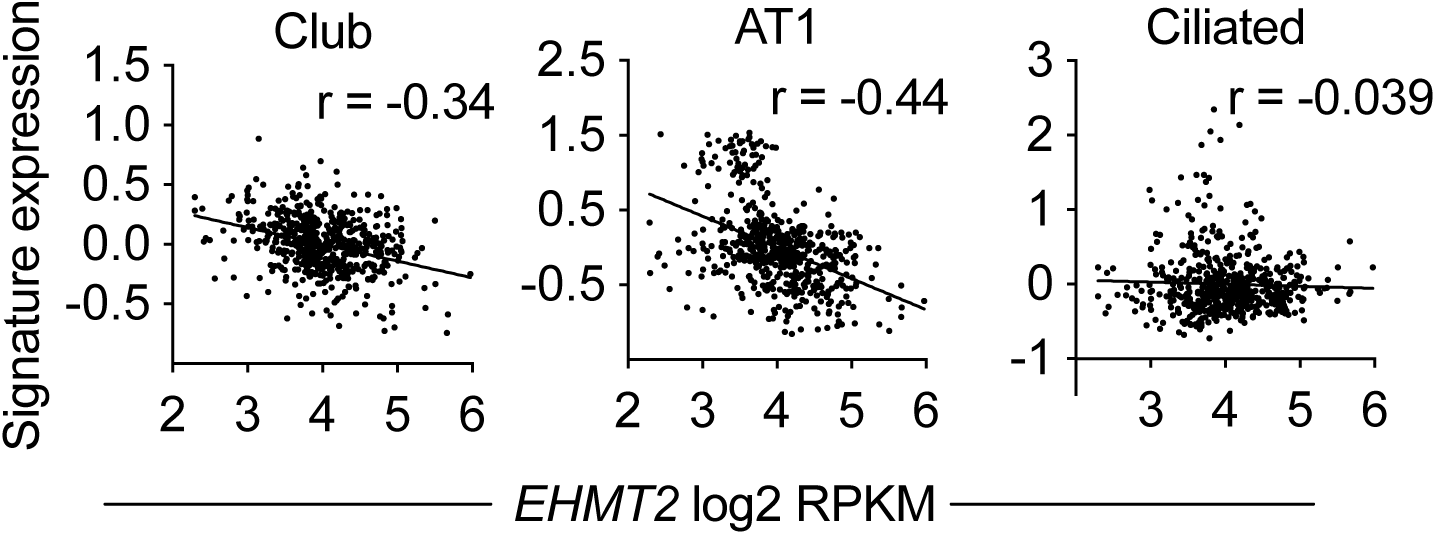
Spearman’s rank correlation analysis between orthogonal human alveolar gene signatures and G9a *EHMT2* transcript in 546 human lung adenocarcinomas (n=546, Linear regression analysis p<0.0001)

**Figure 4–figure supplement 1.**
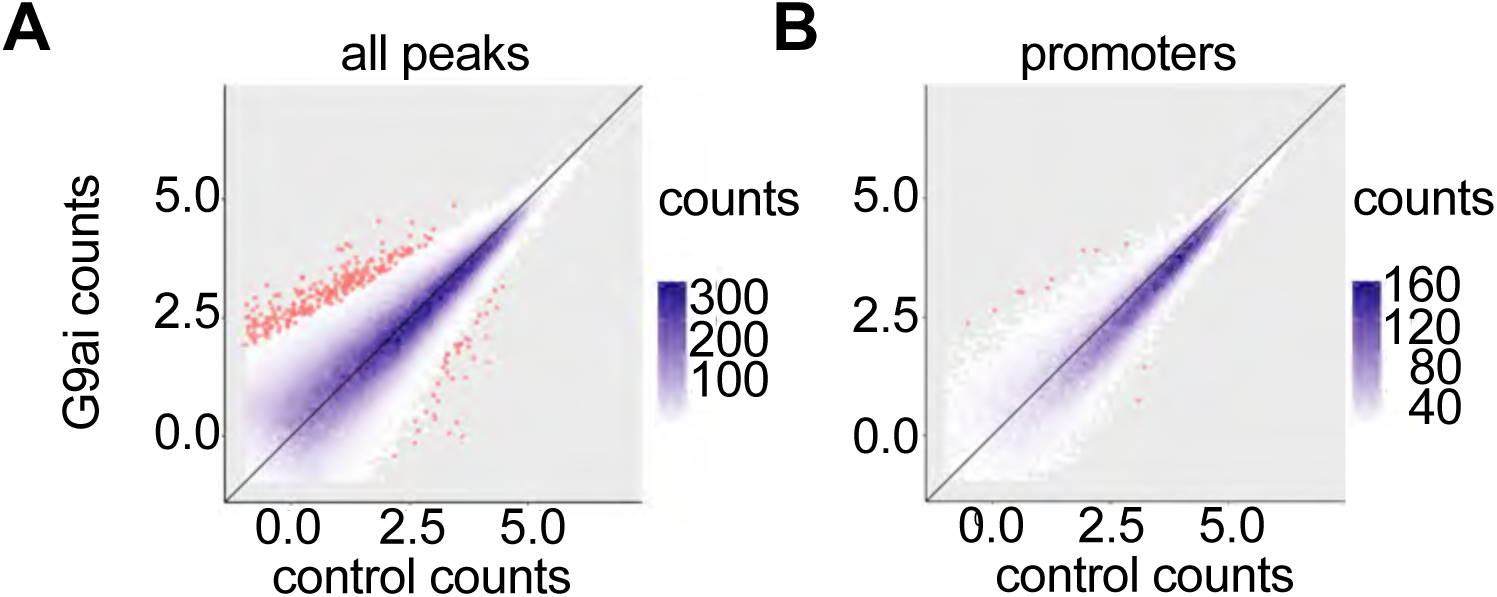
G9a inhibition does not induce broad changes in chromatin accessibility. Scatter plots showing correlation of relative ATAC-seq tag counts for all peaks **(A)** or promoters only **(B)**, from vehicle control (control) and G9a-inhibited (G9ai) tumor-propagating cells (TPCs) isolated from tumorspheres 5 days after inhibitor treatment. Overall 331 peaks and 57 peaks demonstrated increased and decreased accessibility, respectively (a) (n=3; FDR <0.05; fold change>1.5).

**Figure 4–figure supplement 2.**
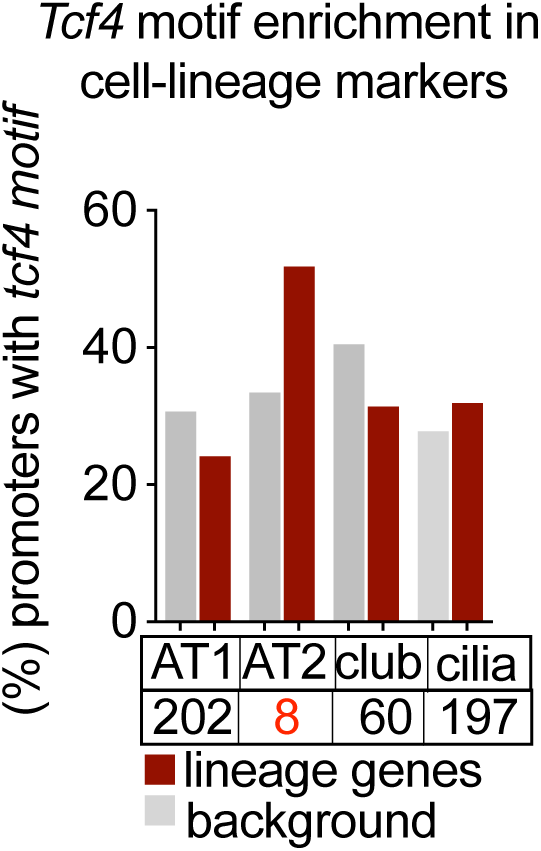
Unbiased enrichment analysis of transcription factor (TF) binding motifs, revealing enrichment of *Tcf4* motifs in promoter regions of distal alveolar cell-lineage gene signatures vs. background, depicted in red and gray color bars, respectively. Bottom table displays how *Tcf4* motifs rank in each of the cell lineage gene signatures.

**Figure 4–figure supplement 3.**
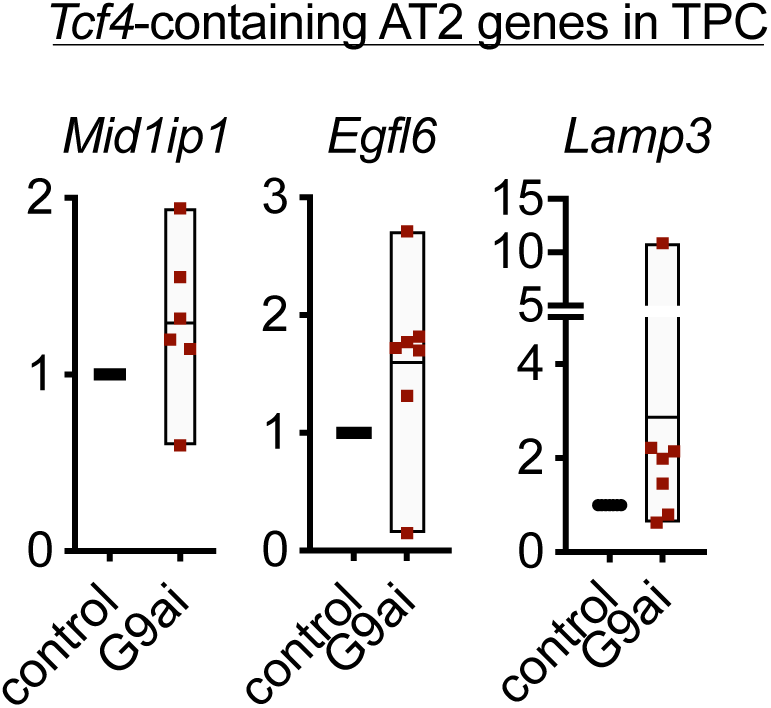
Relative expression of TCF4-containing AT2 markers in TPCs, treated with G9ai vs. control (n=6, mean±SEM, p<0.05).

**Figure 4–figure supplement 4.**
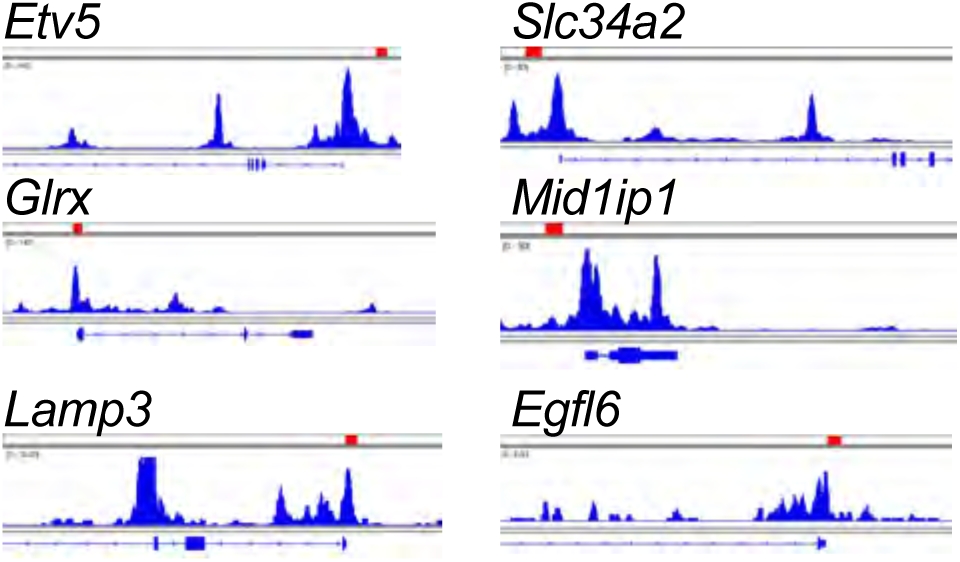
Integrated genome viewer tracks of ATAC-seq generated from TPCs. Tracks showing location of *tcf4* motifs within promoters of AT2-lineage markers (red rectangles, *Tcf4* motif locations).

**Figure 6–figure supplement 1.**
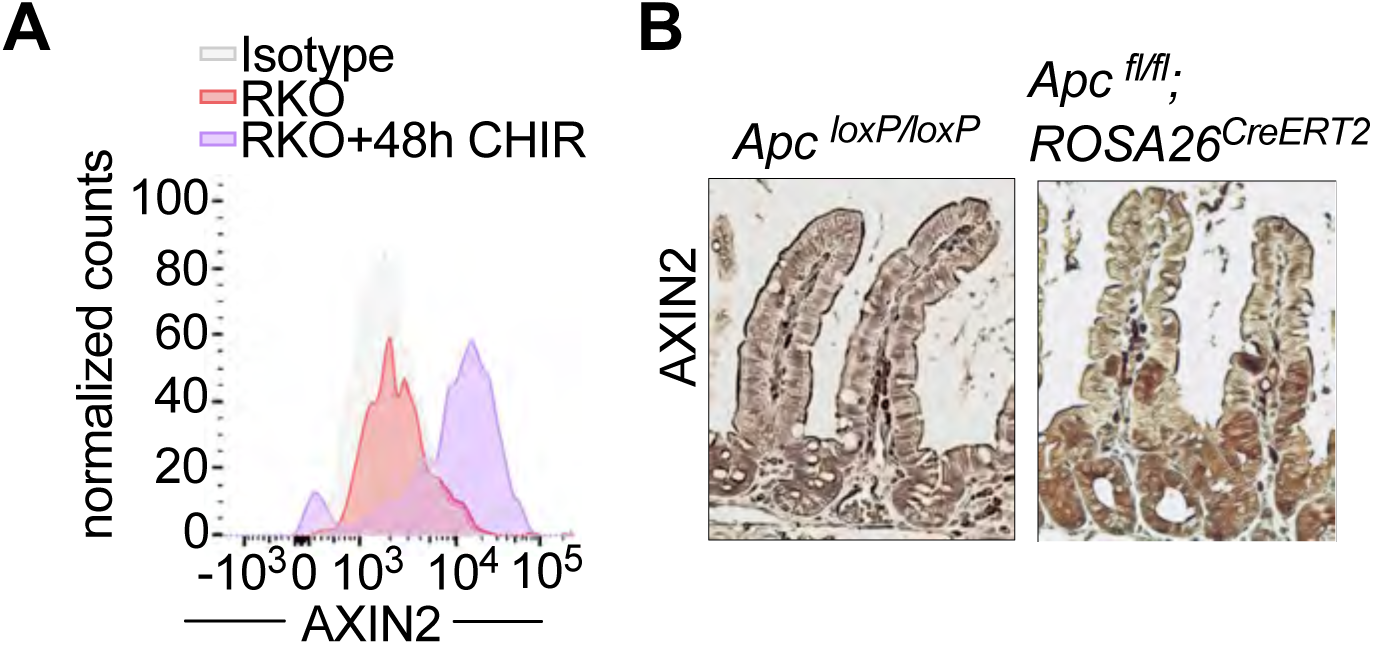
Calibration of Axin2 Antibody. **(A)** Flow cytometry of AXIN2 following 48h of GSK3β inhibitor treatment in the RKO colorectal cell line (RKO + 48h CHIR) compared to vehicle control (RKO) (Isotype depicted in grey; RKO in orange; CHIR-treated RKO in purple). **(B)** Calibration of an Axin2 antibody used in Figure 6A and S6A using immunohistochemistry of AXIN2 in *APC*^fl/fl^ and *APC*^fl/fl^; *ROSA26*^CreERT2^ villi (n=4)

**Figure 6–figure supplement 2.**
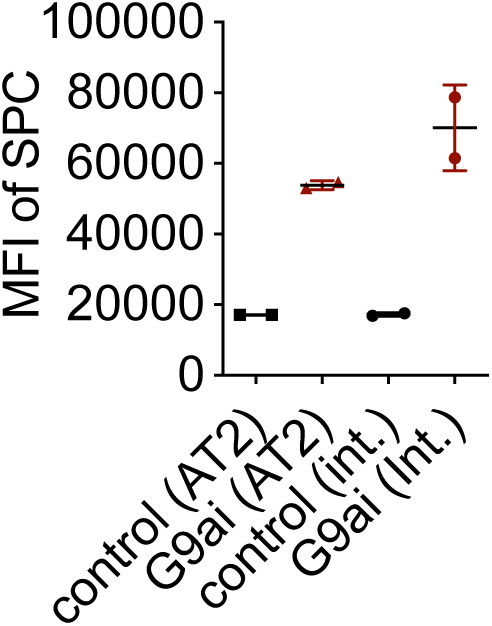
Mean fluorescence intensity (MFI) of SPC in AT2 and intermediate alveosphere-derived cells, following G9a inhibition vs. control (n=2±SEM; two-tailed t-test, p<0.05).

**Figure 6–figure supplement 3.**
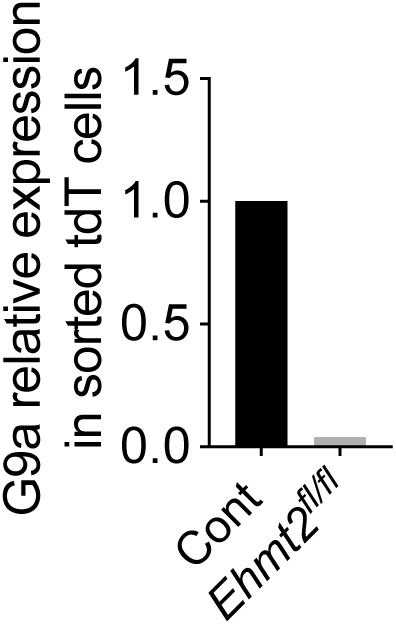
Relative expression of G9a transcript in pooled tdT sorted cells from *Ehmt*^*fl/fl*^ vs. control (n=6)

**Figure 6–figure supplement 4.**
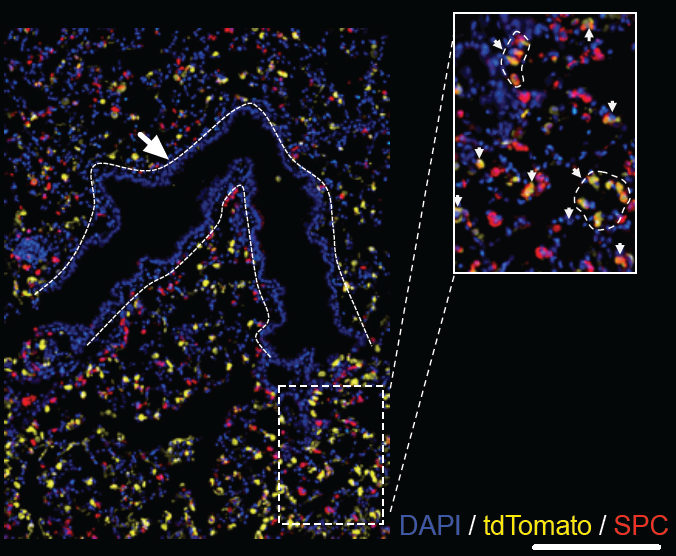
Representative image showing TdTomato-expressing cells a (yellow) and SPC-expressing cells following 5 days of intratracheal infection with *AAV9-Cre*. Dashed lines mark airways. Inset magnification shows colocalization of tdTomato-expressing cells (yellow) and SPC (Red).

**Figure 6–figure supplement 5.**
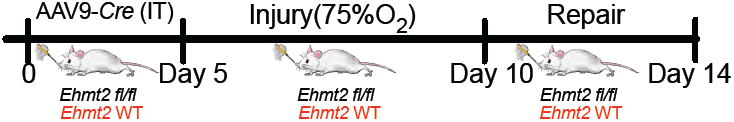
Schematic representation of Hyperoxic (75% O_2_) experiment. WT and *Ehmt2*^*fl/fl*^ mice were infected with *AAV9-Cre*. 5 days following infection, Mice were exposed to Hyperoxic conditions for 5 days and allowed to recover for 4 days.

**Figure 6–figure supplement 6.**
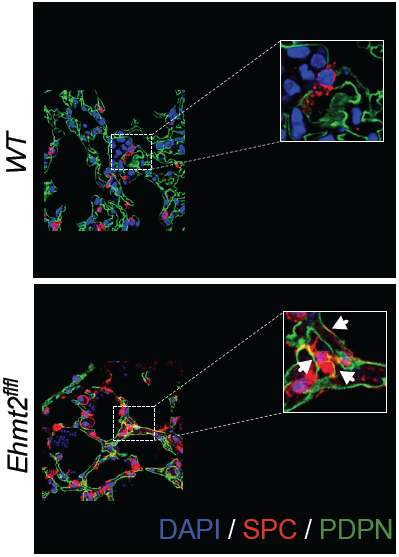
Representative images of SPC+PDPN+ double positive cells. White Arrows in inset show double positive cells in *Ehmt2*^*fl/fl*^ mice or single stained SPC or PDPN – only in WT mice, respectively.

**Figure 6–figure supplement 7.**
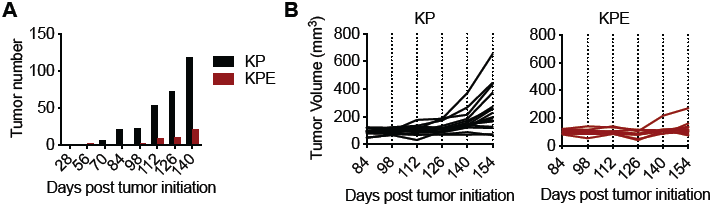
Decreased tumor burden in KPE mice. **(A)** Graph shows cumulative tumor number detected by μ-CT scan at different time points following tumor initiation in KP mice (black) (n=16) and KPE mice (Red) (n=10). **(B)** Graph shows tumor volume quantified by μ-CT scan at different time points following tumor initiation in KP mice (black) and KPE (Red).

## Supplementary Information

**Table S1.**
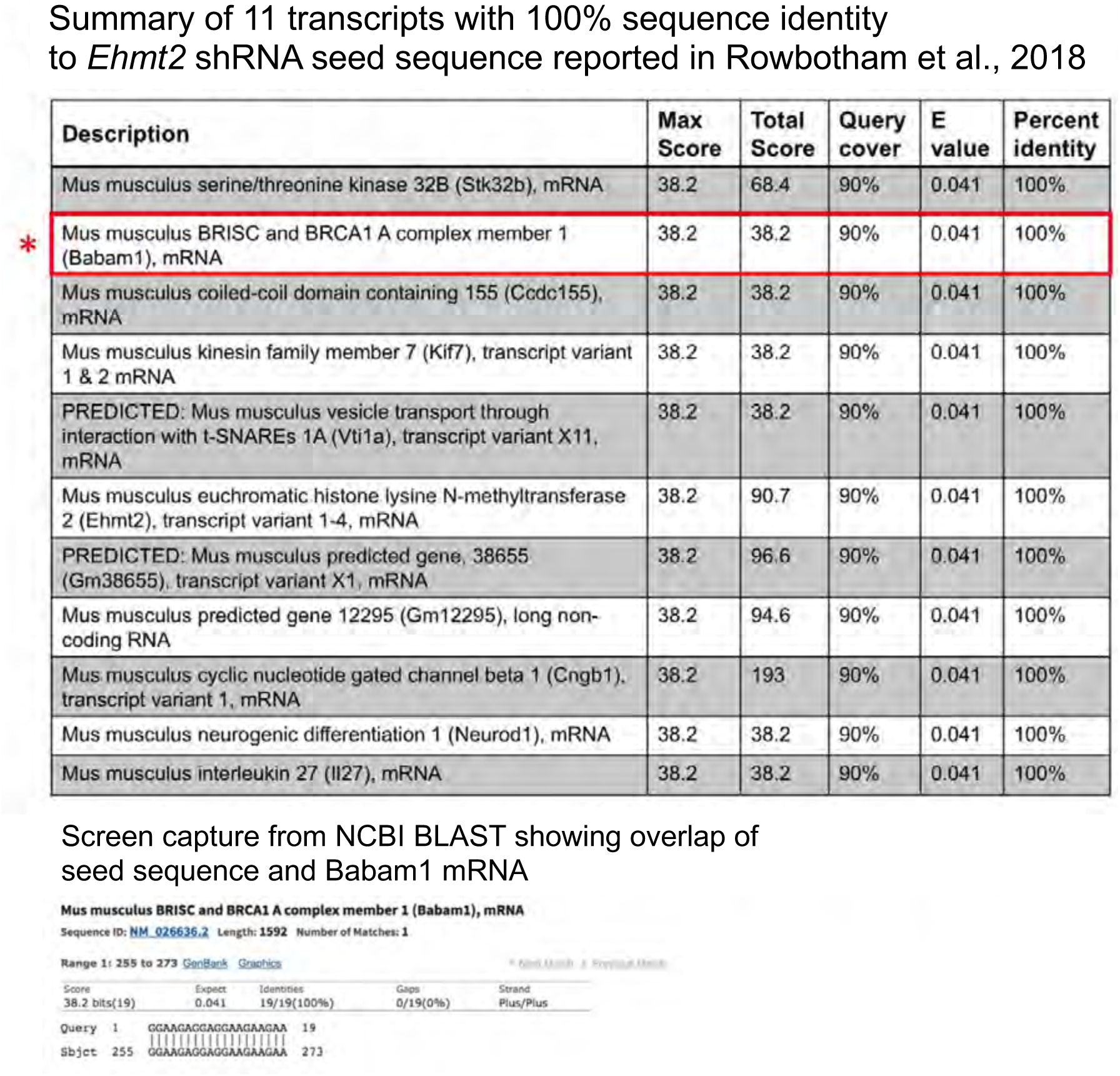

**Table S2.**
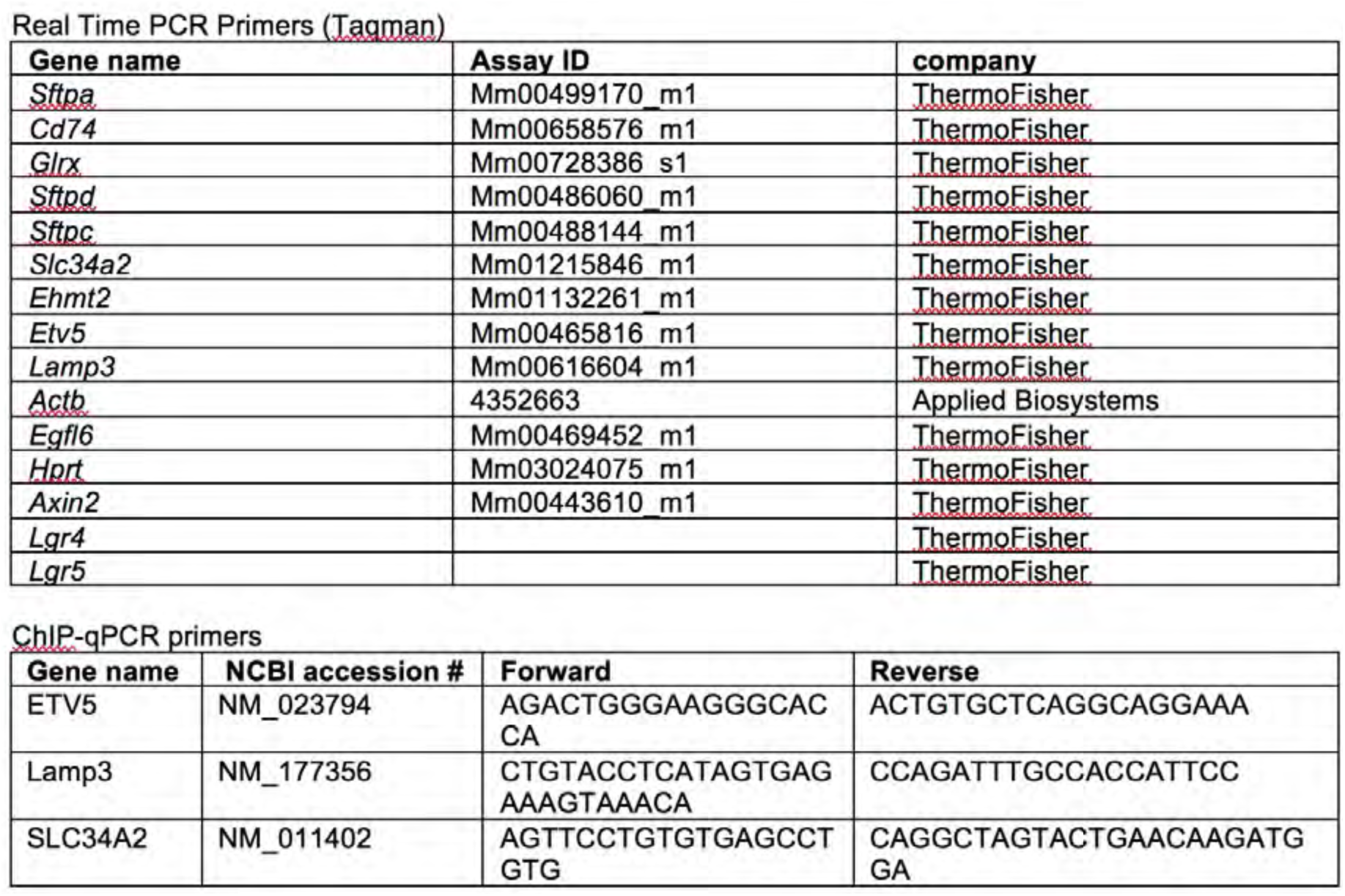

### Supplemental Experimental Procedures

#### Transduction of primary KP tumor cells

shRNAs containing the following *Ehmt2* hairpins shG9.1: 5’ acagcaagtctgaagtcgaa 3’, shG9a.2: 5’ cactgtcaccgtcggcgatga 3’ were synthesized, cloned into a mirE backbone and then subsequently sub-cloned into pInducer-10(Fellmann et al., 2013; Meerbrey et al., 2011) and transfected with packaging constructs into 293T cells using Lipofectamine 2000 (Thermo Fisher). Viral supernatants were collected 72h-post transfection and subsequently concentrated using ultracentrifuge at 25,000 RPM for 2 hours to generate high-titer virus. Freshly-sorted primary KP tumor cells were infected over-night using ultra-low attachment plates (Corning) and subsequently transplanted or grown in Matrigel as tumorspheres.

#### Flow cytometry of TPC and tumorspheres

For TPC sorting and flow cytometry of primary and tumorsphere-derived TPC, single cell preparations were performed as previously described(Zheng et al., 2013) with the following modifications. Immune cell lineage content was excluded by sorting using Biotin-conjugated CD45 (BD, 553078, 30-F11,1:200), CD31(BD, 553371, MEC13.3, 1:200), Ter119 (BD, 553672, Ter119, 1:200), thereafter Biotin-conjugated antibodies were detected with Phycoerythrin (PE)/Cy7 Streptavidin (Biolegend, 405206: 1:300), and subsequently stained for CD24 PerCP-eFluor 710 (eBioscience, 46-0242, M1/69: 1:300), ITGB4-PE (Biolegend, 123602, 346-11A, 1:20) and Notch1 (Biolegend, 130613, HMN1-12, 1:80), Notch2 (Biolegend, 130714, HMN-2-35, 1:80), Notch3 (eBioescience, 17-5763-82, HMN3-133, 1:80), Notch4 (Biolegend, 128413, HMN4-14, 1:80). All anti Notch antibodies are Allophycocyanin (APC) conjugated and used as a pool.

For intracellular staining of sorted TPC and dissociated single-cell preparations from tumorspheres, cells were fixed with 4% PFA for 10 min, washed, permeabilized with 0.25% Triton X-100 for 15 min and then subsequently blocked in 4% BSA containing Fc block (BD, 553141, 2.4/G2, 1:1000) for 30 min. Staining was done in blocking buffer with 0.05% Triton X-100. The following antibodies were used for flow cytometry: CD74-BUV395 (BD, 740274, In-1, 1:25), Pro-SPC (Abcam, ab170699: 1:200), G9a (Abcam, ab185050, EPR18894). Podoplanin (ThermoFisher, MA5-16113, 8.1.1 1:200) For secondary antibody staining, Alexa Fluor 488 antibody (ThermoFisher, A-21206, 1:500) was used. All flow cytometry analysis was performed using FlowJo software.

#### Tumorsphere immunofluorescence (IF)

KP tumorspheres were either stained in Matrigel or were paraffin-embedded, sectioned and stained as previously described(Hegab et al., 2015). For Matrigel preparations, tumorspheres were fixed in 4% PFA for 40 min, washed 3 × 5 min, permeabilized with 0.5% Triton X-100 for 30 min then blocked in 4% BSA for 30-60 min. Staining was performed in blocking buffer with 0.05% Triton X-100 and washes were performed with 0.1% Triton X-100. Tumorspheres were imaged using a Leica SPE confocal microscope. The following antibodies were used for IF: BrdU (NeoMarkers, MS-1058-PO, BRD.3 1:200), cleaved caspase-3 (Cell Signaling Technology, 9661 1:300) pro-SPC (Abcam, ab170699: 1:200), CC10 (SantaCruz, 9772, 1:200) FoxJ1 (eBioscience, 14-9965-82, 2A5, 1:25), RAGE (R&D, 175410, MAB1179, 1:100).

#### BrdU incorporation in tumorspheres

BrdU labeling reagent (ThermoFisher) was incorporated for 3 hours. Tumorspheres were then processed according to IF procedures described in the previous section.

#### Quantification of tumorsphere IF images

Images were taken using SP5 Confocal (Leica). Single projection images of 4-6 z-stack sections from at least 15-20 tumorspheres were constructed and analyzed in the Matlab software package (version R2016b by Mathworks, Natick, MA). Individual cell nuclei were segmented using regional intensity maxima and watershed thresholding on the DAPI channel, and then scored by the presence of relevant IF signal above a global intensity threshold in the area immediately surrounding each nucleus. For BrdU images signal was calculated in each nucleus around intensity maxima. Quality control images were also created by superimposing the cell scoring mask on the raw image data.

#### In vivo treatment with G9a inhibitor UNC0642

6 mice per group were treated with either vehicle (60% PEG400/40% H20) or with UNC0642 10 mg/kg, IP, daily (60% PEG400/40% H_2_0) for a duration of 6 days. Lungs were harvested and sorted for CD24^−^ as previously described(Barkauskas et al., 2013; McQualter et al., 2010)

#### Transmission Electron microscopy (TEM)

Lung tumorspheres were first fixed in modified Karnovsky’s fixative and then post-fixed in freshly prepared 1% aqueous potassium ferrocyanide-osmium tetroxide (EM Sciences, Hatfield, PA), for 2h followed by overnight incubation in 0.5% Uranyl acetate at 4^0^C. The samples were then dehydrated followed by propylene oxide (each step was for 15 min) and embedded in Eponate 12 (Ted Pella, Redding, CA). Ultrathin sections (80 nm) were cut with an Ultracut microtome (Leica), stained with 0.2% lead citrate and examined in a JEOL JEM-1400 transmission electron microscope (TEM) at 80kV. Digital imaged were captured with a GATAN Ultrascan 1000 CCD camera. For each biological replicate, 20 randomly selected cells in each treatment were manually scored and quantified for lamellar bodies. Sections were imaged at low and high magnifications, scale bars 2 µm and 0.5, µm respectively.

#### Western blot for TPCs and tumorspheres

TPCs and non-TPCs were sorted from 60 tumor-bearing mice to generate material for Western. Whole cell extracts were lysed in RIPA buffer (50mM Tris-HCl, 150mM NaCl, 1% Deoxycholate 0.1% SDS 1% Triton-X100) supplemented with Halt protease and phosphatase inhibitor cocktails (ThermoFisher). Lysates were quantified using pierce BCA protein assay kit (Pierce). To detect histones in tumorspheres, acid extraction method was used as previously described(Shechter et al., 2007). Antibodies used for Western blotting of TPCs and nonTPCs: G9a (Abcam, ab185050, EPR18894, 1:1000), Actin (BD, 612656, C4, 1:20,000), was used as loading control. In tumorspheres; H3K9me2 (Abcam, Ab1220, 1:1000), H3K9me3 (Active Motif, 39161 1:1000) Histone H3 (Cell Signaling Technology, 3638, 96C10, 1:1000), was used as loading control.

#### qRT-PCR analysis

RNA isolation for tumorspheres was performed using the RNAeasy Micro plus kit (Qiagen). Tumorspheres were dissociated as described. Samples were then measured using Nanodrop and subsequently reverse transcribed using SuperScript III (ThermoFisher). For TPCs an additional step of pre-amplification was performed using TaqMan Preamp Master Mix (ThermoFisher). qRT-PCR was performed using Fast Advanced PCR Master Mix (ThermoFisher). *Hprt* and *Actin* were used as reference genes for all assays. All Taqman Assays used in the study are described in Table1.

#### Tumorsphere subcellular fractionation

Pooled primary KP tumorspheres (n=6) from G9a-treated and control Matrigel cultures were extracted, fractionated and processed to generate cytoplasmic, nuclear and chromatin fractions, using Subcellular Protein Fractionation Kit Thermo Fisher (78840), as defined by the protocol.

#### Antibody validation assays

Axin2 validation was conducted using both human RKO cells and intestinal sections of *APC*^*loxp/loxp*^ vs. *APC*^*fl/fl*^; *ROSA26*^*CreERT2*^. Briefly, RKO cells were GSK3-β (CHIR)-treated for 48h vs. controls, harvested, 4% paraformaldehyde fixed for 10 min and subsequently washed and permeabilized with 0.25% Triton X-100. Cells were then blocked with 2.5% horse serum and stained for Axin2 Abcam (ab109307, EPR2005, 1:100) for flow cytometry. Intestinal epithelial sections were stained with Axin2 (ab109307, EPR2005, 1:25, AR; Citrate pH=6).

#### RNA-sequencing data for tumorspheres

5 biological repetitions of primary tumorspheres were generated from pooled KP tumors from 6-10 mice per experiment. In each experiment matched KP tumorspheres were treated with either vehicle or G9a inhibitor (UNC0642). Total RNA was extracted using Qiagen RNeasy kit as per the manufacturer’s protocol and quality control of RNA samples was performed to determine their quantity and quality. The concentration of RNA samples was determined using NanoDrop 8000 (Thermo Scientific) and the integrity of RNA was determined by Fragment Analyzer (Advanced Analytical Technologies). 0.5-100 ng of total RNA was used as an input for library preparation using TruSeq RNA Sample Preparation Kit v2 (Illumina). Size of RNA-seq libraries was confirmed using 4200 TapeStation and High Sensitivity D1K screen tape (Agilent Technologies). Library concentration was determined by a qPCR-based method using Library quantification kit (KAPA). The libraries were multiplexed and then sequenced on Illumina HiSeq4000 (Illumina) to generate 30M of single end 50 base pair reads.

RNA sequencing data were analyzed with HTSeqGenie(Pau GB, 2012)in BioConductor(Huber et al., 2015) as follows: first, reads with low nucleotide qualities (70% of bases with quality <23) or rRNA and adapter contamination were removed. Reads were then aligned to the reference genome GRCm38 using GSNAP(Wu and Nacu, 2010). Alignments that were reported by GSNAP as “uniquely mapping” were used for subsequent analysis. Gene expression levels were quantified as Reads Per Kilobase of exon model per Million mapped reads normalized by size factor (nRPKM), defined as number of reads aligning to a gene in a sample / (total number of uniquely mapped reads for that sample × gene length × size factor).

#### Cancer Genome Atlas (TCGA) RNA-seq data analysis

RNA-sequencing data for 546 lung adenocarcinoma tumors from TCGA(Cancer Genome Atlas Research, 2014) were obtained from the National Cancer Institute Genomic Data Commons (https://gdc.cancer.gov). We employed the same approach for RNAseq data processing and quantification, with human reference genome GRCh38.

#### Motif enrichment near lineage markers

For each cell lineage, function findMotifs.pl from HOMER 4.7(Heinz et al., 2010) was used to identify enriched motifs near cell lineage markers from (Treutlein et al., 2014). Motif enrichment was performed with promoter regions defined from −2000bp to 500bp relative to TSS, and with the background sets defined as the full genome. Locations of the TCF4 motif near cell lineage markers (from −2000bp to 500bp relative to TSS) were determined using findMotifs.pl from HOMER 4.7. The TCF4 motif file was obtained from the HomerMotifDB (http://homer.ucsd.edu/homer/motif/HomerMotifDB/homerResults.html).

#### Code and data availability

A source code used in this study has been made available in the R computer language. This package is freely available under the Creative Commons 3.0 license and can be found at https://github.com/anneleendaemen/G9a.CellIdentity.Lung

## References

Balis, J.U., and Conen, P.E. (1964). The Role of Alveolar Inclusion Bodies in the Developing Lung. Lab Invest 13, 1215–1229.

Barkauskas, C.E., Cronce, M.J., Rackley, C.R., Bowie, E.J., Keene, D.R., Stripp, B.R., Randell, S.H., Noble, P.W., and Hogan, B.L. (2013). Type 2 alveolar cells are stem cells in adult lung. J Clin Invest 123, 3025–3036.

Batlle, E., and Clevers, H. (2017). Cancer stem cells revisited. Nat Med 23, 1124–1134.

Bauer, A., Chauvet, S., Huber, O., Usseglio, F., Rothbacher, U., Aragnol, D., Kemler, R., and Pradel, J. (2000). Pontin52 and reptin52 function as antagonistic regulators of beta-catenin signalling activity. EMBO J 19, 6121–6130.

Beck, B., and Blanpain, C. (2013). Unravelling cancer stem cell potential. Nat Rev Cancer 13, 727–738.

Buenrostro, J.D., Giresi, P.G., Zaba, L.C., Chang, H.Y., and Greenleaf, W.J. (2013). Transposition of native chromatin for fast and sensitive epigenomic profiling of open chromatin, DNA-binding proteins and nucleosome position. Nat Methods 10, 1213–1218.

Cancer Genome Atlas Research, N. (2014). Comprehensive molecular profiling of lung adenocarcinoma. Nature 511, 543–550.

Chen, M.W., Hua, K.T., Kao, H.J., Chi, C.C., Wei, L.H., Johansson, G., Shiah, S.G., Chen, P.S., Jeng, Y.M., Cheng, T.Y., et al. (2010). H3K9 histone methyltransferase G9a promotes lung cancer invasion and metastasis by silencing the cell adhesion molecule Ep-CAM. Cancer Res 70, 7830–7840.

Chen, X., Skutt-Kakaria, K., Davison, J., Ou, Y.L., Choi, E., Malik, P., Loeb, K., Wood, B., Georges, G., Torok-Storb, B., et al. (2012). G9a/GLP-dependent histone H3K9me2 patterning during human hematopoietic stem cell lineage commitment. Genes Dev 26, 2499–2511.

Collins, R., and Cheng, X. (2010). A case study in cross-talk: the histone lysine methyltransferases G9a and GLP. Nucleic Acids Res 38, 3503–3511.

de Lau, W., Barker, N., Low, T.Y., Koo, B.K., Li, V.S., Teunissen, H., Kujala, P., Haegebarth, A., Peters, P.J., van de Wetering, M., et al. (2011). Lgr5 homologues associate with Wnt receptors and mediate R-spondin signalling. Nature 476, 293–297.

Desai, T.J., Brownfield, D.G., and Krasnow, M.A. (2014). Alveolar progenitor and stem cells in lung development, renewal and cancer. Nature 507, 190–194.

Domen, J., and Weissman, I.L. (1999). Self-renewal, differentiation or death: regulation and manipulation of hematopoietic stem cell fate. Mol Med Today 5, 201–208.

Easwaran, H., Tsai, H.C., and Baylin, S.B. (2014). Cancer epigenetics: tumor heterogeneity, plasticity of stem-like states, and drug resistance. Mol Cell 54, 716–727.

Epsztejn-Litman, S., Feldman, N., Abu-Remaileh, M., Shufaro, Y., Gerson, A., Ueda, J., Deplus, R., Fuks, F., Shinkai, Y., Cedar, H., et al. (2008). De novo DNA methylation promoted by G9a prevents reprogramming of embryonically silenced genes. Nat Struct Mol Biol 15, 1176–1183.

Frank, D.B., Peng, T., Zepp, J.A., Snitow, M., Vincent, T.L., Penkala, I.J., Cui, Z., Herriges, M.J., Morley, M.P., Zhou, S., et al. (2016). Emergence of a Wave of Wnt Signaling that Regulates Lung Alveologenesis by Controlling Epithelial Self-Renewal and Differentiation. Cell Rep 17, 2312–2325.

Hemberger, M., Dean, W., and Reik, W. (2009). Epigenetic dynamics of stem cells and cell lineage commitment: digging Waddington’s canal. Nat Rev Mol Cell Biol 10, 526–537.

Huang, T., Zhang, P., Li, W., Zhao, T., Zhang, Z., Chen, S., Yang, Y., Feng, Y., Li, F., Shirley Liu, X., et al. (2017). G9A promotes tumor cell growth and invasion by silencing CASP1 in non-small-cell lung cancer cells. Cell Death Dis 8, e2726.

Jackson, E.L., Willis, N., Mercer, K., Bronson, R.T., Crowley, D., Montoya, R., Jacks, T., and Tuveson, D.A. (2001). Analysis of lung tumor initiation and progression using conditional expression of oncogenic K-ras. Genes Dev 15, 3243–3248.

Jonkers, J., Meuwissen, R., van der Gulden, H., Peterse, H., van der Valk, M., and Berns, A. (2001). Synergistic tumor suppressor activity of BRCA2 and p53 in a conditional mouse model for breast cancer. Nat Genet 29, 418–425.

Kim, Y., Lee, H.M., Xiong, Y., Sciaky, N., Hulbert, S.W., Cao, X., Everitt, J.I., Jin, J., Roth, B.L., and Jiang, Y.H. (2017). Targeting the histone methyltransferase G9a activates imprinted genes and improves survival of a mouse model of Prader-Willi syndrome. Nat Med 23, 213–222.

Lee, C.L., Moding, E.J., Huang, X., Li, Y., Woodlief, L.Z., Rodrigues, R.C., Ma, Y., and Kirsch, D.G. (2012). Generation of primary tumors with Flp recombinase in FRT-flanked p53 mice. Dis Model Mech 5, 397–402.

Lee, J.H., Kim, J., Gludish, D., Roach, R.R., Saunders, A.H., Barrios, J., Woo, A.J., Chen, H., Conner, D.A., Fujiwara, Y., et al. (2013). Surfactant protein-C chromatin-bound green fluorescence protein reporter mice reveal heterogeneity of surfactant protein C-expressing lung cells. Am J Respir Cell Mol Biol 48, 288–298.

Lee, J.S., Kim, Y., Kim, I.S., Kim, B., Choi, H.J., Lee, J.M., Shin, H.J., Kim, J.H., Kim, J.Y., Seo, S.B., et al. (2010). Negative regulation of hypoxic responses via induced Reptin methylation. Mol Cell 39, 71–85.

Liberzon, A., Birger, C., Thorvaldsdottir, H., Ghandi, M., Mesirov, J.P., and Tamayo, P. (2015). The Molecular Signatures Database (MSigDB) hallmark gene set collection. Cell Syst 1, 417–425.

Liu, F., Barsyte-Lovejoy, D., Li, F., Xiong, Y., Korboukh, V., Huang, X.P., Allali-Hassani, A., Janzen, W.P., Roth, B.L., Frye, S.V., et al. (2013). Discovery of an in vivo chemical probe of the lysine methyltransferases G9a and GLP. J Med Chem 56, 8931–8942.

Madisen, L., Zwingman, T.A., Sunkin, S.M., Oh, S.W., Zariwala, H.A., Gu, H., Ng, L.L., Palmiter, R.D., Hawrylycz, M.J., Jones, A.R., et al. (2010). A robust and high-throughput Cre reporting and characterization system for the whole mouse brain. Nat Neurosci 13, 133–140.

Mainardi, S., Mijimolle, N., Francoz, S., Vicente-Duenas, C., Sanchez-Garcia, I., and Barbacid, M. (2014). Identification of cancer initiating cells in K-Ras driven lung adenocarcinoma. Proc Natl Acad Sci U S A 111, 255–260.

Mao, Y.Q., and Houry, W.A. (2017). The Role of Pontin and Reptin in Cellular Physiology and Cancer Etiology. Front Mol Biosci 4, 58.

McQualter, J.L., Yuen, K., Williams, B., and Bertoncello, I. (2010). Evidence of an epithelial stem/progenitor cell hierarchy in the adult mouse lung. Proc Natl Acad Sci U S A 107, 1414–1419.

Nabhan, A.N., Brownfield, D.G., Harbury, P.B., Krasnow, M.A., and Desai, T.J. (2018). Single-cell Wnt signaling niches maintain stemness of alveolar type 2 cells. Science 359, 1118–1123.

Pacheco-Pinedo, E.C., Durham, A.C., Stewart, K.M., Goss, A.M., Lu, M.M., Demayo, F.J., and Morrisey, E.E. (2011). Wnt/beta-catenin signaling accelerates mouse lung tumorigenesis by imposing an embryonic distal progenitor phenotype on lung epithelium. J Clin Invest 121, 1935–1945.

Reya, T., Morrison, S.J., Clarke, M.F., and Weissman, I.L. (2001). Stem cells, cancer, and cancer stem cells. Nature 414, 105–111.

Robinson, J.T., Thorvaldsdottir, H., Winckler, W., Guttman, M., Lander, E.S., Getz, G., and Mesirov, J.P. (2011). Integrative genomics viewer. Nat Biotechnol 29, 24–26.

Robinson, M.D., and Oshlack, A. (2010). A scaling normalization method for differential expression analysis of RNA-seq data. Genome Biol 11, R25.

Ruijtenberg, S., and van den Heuvel, S. (2016). Coordinating cell proliferation and differentiation: Antagonism between cell cycle regulators and cell type-specific gene expression. Cell Cycle 15, 196–212.

Shackleton, M., Quintana, E., Fearon, E.R., and Morrison, S.J. (2009). Heterogeneity in cancer: cancer stem cells versus clonal evolution. Cell 138, 822–829.

Shibata, H., Komura, S., Yamada, Y., Sankoda, N., Tanaka, A., Ukai, T., Kabata, M., Sakurai, S., Kuze, B., Woltjen, K., et al. (2018). In vivo reprogramming drives Kras-induced cancer development. Nat Commun 9, 2081.

Shinkai, Y., and Tachibana, M. (2011). H3K9 methyltransferase G9a and the related molecule GLP. Genes Dev 25, 781–788.

Sutherland, K.D., Song, J.Y., Kwon, M.C., Proost, N., Zevenhoven, J., and Berns, A. (2014). Multiple cells-of-origin of mutant K-Ras-induced mouse lung adenocarcinoma. Proc Natl Acad Sci U S A 111, 4952–4957.

Tammela, T., Sanchez-Rivera, F.J., Cetinbas, N.M., Wu, K., Joshi, N.S., Helenius, K., Park, Y., Azimi, R., Kerper, N.R., Wesselhoeft, R.A., et al. (2017). A Wnt-producing niche drives proliferative potential and progression in lung adenocarcinoma. Nature 545, 355–359.

Treutlein, B., Brownfield, D.G., Wu, A.R., Neff, N.F., Mantalas, G.L., Espinoza, F.H., Desai, T.J., Krasnow, M.A., and Quake, S.R. (2014). Reconstructing lineage hierarchies of the distal lung epithelium using single-cell RNA-seq. Nature 509, 371–375.

Widschwendter, M., Fiegl, H., Egle, D., Mueller-Holzner, E., Spizzo, G., Marth, C., Weisenberger, D.J., Campan, M., Young, J., Jacobs, I., et al. (2007). Epigenetic stem cell signature in cancer. Nat Genet 39, 157–158.

Wu, T.D., and Nacu, S. (2010). Fast and SNP-tolerant detection of complex variants and splicing in short reads. Bioinformatics 26, 873–881.

http://www.cancer.org (2019). https://www.cancer.org/cancer/non-small-cell-lung-cancer/about/key-statistics.html.

Xu, X., Rock, J.R., Lu, Y., Futtner, C., Schwab, B., Guinney, J., Hogan, B.L., and Onaitis, M.W. (2012). Evidence for type II cells as cells of origin of K-Ras-induced distal lung adenocarcinoma. Proc Natl Acad Sci U S A 109, 4910–4915.

Zacharias, W.J., Frank, D.B., Zepp, J.A., Morley, M.P., Alkhaleel, F.A., Kong, J., Zhou, S., Cantu, E., and Morrisey, E.E. (2018). Regeneration of the lung alveolus by an evolutionarily conserved epithelial progenitor. Nature 555, 251–255.

Zepp, J.A., Zacharias, W.J., Frank, D.B., Cavanaugh, C.A., Zhou, S., Morley, M.P., and Morrisey, E.E. (2017). Distinct Mesenchymal Lineages and Niches Promote Epithelial Self-Renewal and Myofibrogenesis in the Lung. Cell 170, 1134–1148 e1110.

Zhang, Y., Liu, T., Meyer, C.A., Eeckhoute, J., Johnson, D.S., Bernstein, B.E., Nusbaum, C., Myers, R.M., Brown, M., Li, W., et al. (2008). Model-based analysis of ChIP-Seq (MACS). Genome Biol 9, R137.

Zheng, Y., de la Cruz, C.C., Sayles, L.C., Alleyne-Chin, C., Vaka, D., Knaak, T.D., Bigos, M., Xu, Y., Hoang, C.D., Shrager, J.B., et al. (2013). A rare population of CD24(+)ITGB4(+)Notch(hi) cells drives tumor propagation in NSCLC and requires Notch3 for self-renewal. Cancer Cell 24, 59–74.

Zylicz, J.J., Borensztein, M., Wong, F.C., Huang, Y., Lee, C., Dietmann, S., and Surani, M.A. (2018). G9a regulates temporal preimplantation developmental program and lineage segregation in blastocyst. Elife 7.

## Supplemental References

Fellmann, C., Hoffmann, T., Sridhar, V., Hopfgartner, B., Muhar, M., Roth, M., Lai, D.Y., Barbosa, I.A., Kwon, J.S., Guan, Y., et al. (2013). An optimized microRNA backbone for effective single-copy RNAi. Cell Rep 5, 1704–1713.

Heinz, S., Benner, C., Spann, N., Bertolino, E., Lin, Y.C., Laslo, P., Cheng, J.X., Murre, C., Singh, H., and Glass, C.K. (2010). Simple combinations of lineage-determining transcription factors prime cis-regulatory elements required for macrophage and B cell identities. Mol Cell 38, 576–589.

Huber, W., Carey, V.J., Gentleman, R., Anders, S., Carlson, M., Carvalho, B.S., Bravo, H.C., Davis, S., Gatto, L., Girke, T., et al. (2015). Orchestrating high-throughput genomic analysis with Bioconductor. Nat Methods 12, 115–121.

Meerbrey, K.L., Hu, G., Kessler, J.D., Roarty, K., Li, M.Z., Fang, J.E., Herschkowitz, J.I., Burrows, A.E., Ciccia, A., Sun, T., et al. (2011). The pINDUCER lentiviral toolkit for inducible RNA interference in vitro and in vivo. Proc Natl Acad Sci U S A 108, 3665–3670.

Pau GB, R.C., Lawrence J, Degenhardt M, Wu J, Huntley T, Brauer MM (2012). HTSeqGenie: a software package to analyse high-throughput sequencing experiments.. Bioconductor package 1–12.

